# Expansion of the neocortex and protection from neurodegeneration by *in vivo* transient reprogramming

**DOI:** 10.1101/2023.11.27.568858

**Authors:** Y Shen, S Zaballa, X Bech, A Sancho-Balsells, C Díaz-Cifuentes, G Seyit-Bremer, I Ballasch, N Alcázar, J Alberch, M Abad, M Serrano, R Klein, A Giralt, D del Toro

**Author notes:** co-first (Y Shen, S Zaballa).

## Abstract

Yamanaka factors (YFs) can reverse some aging features in mammalian tissues, but their effects on the brain remain largely unexplored. Here, we induced YFs in the mouse brain in a controlled spatio-temporal manner in two different scenarios: brain development, and adult stages in the context of neurodegeneration. Embryonic induction of YFs perturbed cell identity of both progenitors and neurons, but transient and low-level expression is tolerated by these cells during development. Under these conditions, YFs induction led to expanded neurogenesis, increased number of upper cortical neurons, and enhanced motor and social behavior of adult mice. Additionally, controlled YF induction is tolerated by principal neurons in the adult dorsal hippocampus and prevented the development of several hallmarks of Alzheimer’s disease, including cognitive decline and altered molecular signatures, in the 5xFAD mouse model. Overall, these results highlight the powerful impact of YFs on neurogenesis and their potential use in brain disorders.

**Highlights:**

- Transient Yamanaka factor (YF) expression during development expands neocortex
- YF-treated mice show enhanced cognitive skills
- Intermittent YF expression is tolerated by adult principal hippocampal neurons
- Long-term intermittent YF reprogramming is protective in an AD mouse model

## Introduction

The Yamanaka factors, a quartet of transcription factors comprising Oct4, Sox2, Klf4 and c-Myc (hereafter referred to as YFs), have delineated a pivotal role in cellular biology due to their unparalleled ability to reprogram somatic cells back to a pluripotent state^1^. This discovery has opened new avenues for research, unveiling their potential in tissue rejuvenation^2,3^, reversal of epigenetic modifications and markers of cell damage^4–6^, thus ameliorating phenotypes associated with cellular aging^2,7,8^. Indeed, YFs have been employed to reverse age-associated hallmarks in various peripheral tissues, resulting in improved regeneration of muscle, optic nerve, cardiomyocytes, skin and liver^5,9–13^. Despite these findings, our understanding of their role and broader implications for the nervous system is still in its nascent stages.

Ectopic expression of YFs can induce dedifferentiation and pluripotency of somatic cells under both *in vitro* and *in vivo* conditions, erasing their epigenetic state and thereby their cellular identity^6,14^. This process has been molecularly coupled to induced pluripotent cell (iPSC) formation mechanisms^15,16^. Consequently, continuous expression of these factors in wild-type mice often results in the development of tumoral masses, predominantly teratomas, and leads to mortality within a matter of weeks^17^. Seminal work has confirmed that it is possible to safely express YFs *in vivo* through partial or transient activation^2,9^. When applied to various peripheral tissues, this intermittent activation not only prevents the loss of cell identity but also enhances regeneration without causing cancer^2,4^. Moreover, transient YFs activation has been shown to increase the proliferation potential of multiple tissues through distinct mechanisms: (1) by remodeling the stem cell niche to activate stem proliferation in muscles^11^, and (2) by inducing transient de-differentiation of cardiomyocites to restore their regenerative capability^12,13^ . It remains an open question whether transient YFs induction could enhance proliferation and/or rejuvenation in the nervous system.

Aging stands as the primary risk factor for Alzheimer’s disease (AD), a phenomenon not entirely accounted for by the amyloid hypothesis^18^. Notably, AD, together with most neurodegenerative disorders, presents signatures of “accelerated” aging, including increased oxidation, diminished synaptic plasticity, and reduced metabolism^19–21^. In such scenarios, neurons progressively lose their regenerative capacity during maturation, attributed to alterations in the transcriptomic and chromatin landscapes^22^. From a translational standpoint, YFs have been employed in cell replacement therapies for neurodegenerative diseases, due to their ability to generate iPSCs that can be differentiated to any cell fate^23^. For instance, in Parkinson disease, with the transplantation of dopaminergic neurons derived from YF-induced iPSCs^24^. Although this approach yielded some motor improvements, there were also adverse effects with the growth of teratomas in mice with a severe combined immunodeficiency^25^ . While most reprogramming strategies aim to generate neurons to replace damaged ones, two recent studies have shown that partial reprogramming with YFs can induce epigenetic modifications in neurons and promote axon regeneration after injury^5,26^. However, the application of YFs in mature neurons within the context of neurodegeneration remains unexplored.

Here, we employed a controlled spatio-temporal induction of YFs in the mouse brain across two distinct scenarios: during brain development, and in adult stages within the context of neurodegeneration. Our focus on the impact of YFs on neurogenesis during development was influenced by recent findings that a subset of these factors is endogenously expressed in various neural progenitors at early stages of this phase^27^, and thus their induction during development appears less artificial than initially presumed. Here we report that transient, low-level expression, of YFs amplified neurogenesis resulting in an augmented output of neurons and an enlarged neocortex. This expansion was functionally reflected in enhanced motor and social behavior in adult mice. Since this induction protocol enhanced cognitive skills, we hypothesized that it could exert a similar effect in the context of a neurodegenerative disorder. Thus, we expressed YFs only in mature hippocampal neurons using the 5xFAD mouse model of AD. We show that these neurons tolerate intermittent YFs expression while preserving their cellular identity. In tandem with the safety of our approach, 5xFAD mice with hippocampal YFs expression displayed a rescue of several cognitive impairments. This amelioration was associated with a reduction of several AD-associated phenotypes, including Aβ accumulation, synaptic loss, and proteomic disease profile. Our results establish transient YFs induction as a powerful tool for modulating neurogenesis and may be used to open new therapeutic strategies for brain disorders.

## Results

### Transient YFs induction perturbs cell identity and increases neurogenesis

To study the effects of YFs induction during development, we used the conditional inducible-four factors (i4F) mouse line that allows the induction of YFs in a Tet-ON system^17^. We established an *in vivo* transient reprogramming protocol for systemic expression of YFs by using mice carrying a single copy of the i4F polycistronic cassette and a reverse tetracycline transactivator (rtTA) within the ubiquitously-expressed *Rosa*26 locus (i4F-Rosa)^17^ (Fig. 1A). We performed YFs induction by administering doxycycline (Dox) in the drinking water of pregnant females for 4d (E10.5 to E14.5), using three different concentrations: High (1mg/ml), medium (0.5mg/ml) and low (0.2mg/ml) (Fig. 1A). These concentrations allowed for the assessment of the Dox-dependent effects on the induction of YFs during development (Fig. 1A). We first treated i4F-Rosa embryos and their littermate controls (WT) with high Dox for 4d. We observed that strong YF induction resulted in bigger brains (Movie S1) with a dramatic elongation of the germinal layer at E15.5. This layer was also thinner but with multiple invaginations of the ventricular surface (Fig. 1B-C). Immunostaining for Pax6 (a forebrain progenitor marker) and Sox2 (a neuronal progenitor marker and one of the YFs), showed massive expansion in the number of apical progenitors (Fig. 1D). Despite the increased abundance of these cells, the intensity of Pax6 at the single cell level was reduced (Fig. 1E), suggesting that strong YFs induction during development can perturb cell identity, as has been found in other systems^17,28^. This is evidenced by the loss of identity markers such as Pax6 for progenitors and Ctip2 for neurons, along with the emergence of Nanog expression (Fig. S1A-C). Given that the survival of i4F-Rosa embryos was compromised due to impaired liver hematopoiesis (data not shown), we reduced the concentration of Dox during the 4d induction period.

**Figure 1.**
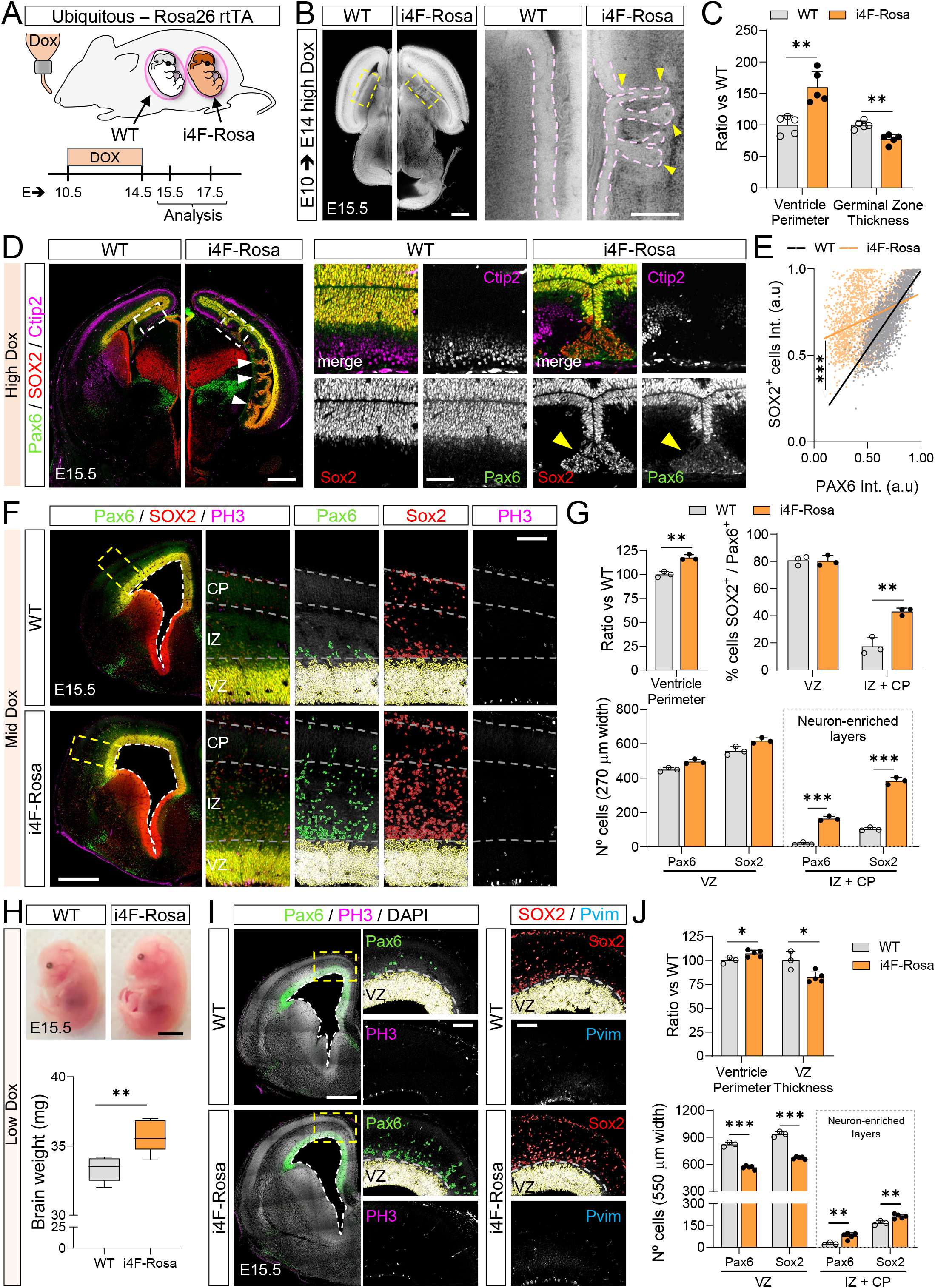
Transient YFs induction perturbs cell identity and increases neurogenesis. (A) Scheme of Dox treatment in wild-type (WT) and i4F-Rosa littermate embryos used in downstream analysis. (B) Sagittal sections of cleared whole-mount brains (E15.5) stained with propidium iodine after high Dox treatment (from E10.5 to E14.5, 4d). Dashed rectangles are shown with higher magnification on the right. Pink dashed lines indicate the germinal zone and yellow arrowheads the presence of invaginations. (C) Quantification of data shown in B. n=5 mice/group. **p<0.01, unpaired Student’s t test. (D) E15.5 coronal brain sections, after 4d of high Dox, were labeled for neuronal progenitors (Pax6, green and Sox2, red) and neurons (Ctip2, magenta). Areas in dashed rectangles are shown with higher magnification on the right. (E) Scatter plot quantification of Pax6 intensity levels of Sox2+ cells. n= 2339 (WT) and 2031 (i4F-Rosa) cells from 3-4 brains/group. ***p<0.001, linear regression slopes, unpaired Student’s t test. (F) Similar experiment as shown in panel D except for E15.5 sections treated with 4d Mid Dox and including mitotic cell marker (PH3, magenta). (G) Quantification of data shown in F. n=3 mice/group. **p<0.01, ***p<0.001, unpaired Student’s t test. (H) E15.5 embryos after 4d Low Dox treatment (from E10.5 to E14.5). Quantification of their brain weight is shown below. n=5-6 mice/group. **p<0.01, unpaired Student’s t test. (I) Similar experiment as shown in panel D and F, except for E15.5 sections treated with 4d Low Dox and including dividing RG cells (Pvim, cyan). Yellow dashed rectangles are shown with higher magnification on the right. Ventricle perimeter is shown with white dashed lines. (J) Quantification of data shown in (I).n=3-5 mice/group. *p<0.05, **p<0.01, ***p<0.001, unpaired Student’s t test. The data are represented as mean ± SEM. Circles indicate values of individual brains or cells (E). Scale bars, 500um (B, D, F, I), 250um inset (B), 100um inset (D,F, I) and 3mm (H).

Medium YFs induction also resulted in an enlarged ventricular zone but without invaginations and loss of Pax6 expression from apical progenitors (Fig. 1F). Notably, the abundance of individual and double-positive Pax6+ and Sox2+ progenitors was significantly higher in neuron-enriched layers (intermediate zone [IZ] and cortical plate [CP]) (Fig. 1G). Similar to high Dox treatment, very few i4F-Rosa embryos reached E17.5 stages, due to altered hematopoiesis.

Next, i4F-Rosa embryos were subjected to low Dox treatment for 4d from E10.5 to E14.5. Remarkably, this condition led to completely viable and bigger embryos with increased brain weight (Fig. 1H). Histological analysis showed increased perimeter of the ventricles that was accompanied by a thinner germinal zone (Fig. 1I-J). The expression of Pax6 in apical progenitors was preserved and there was a significant increase of Pax6+ and Sox2+ progenitors in neuronal enriched-layers (IZ+CP), albeit with lower numbers compared with medium Dox treatment (Fig. 1J). Similar to medium YFs induction, we did not observe changes in the density of cells stained for phosphorylated forms of vimentin (Pvim) and histone H3 (PH3) that label dividing radial glial (RG) and mitotic cells, respectively at E15.5 (Fig. S1D). These results indicate that YFs induction can perturb cell identity and increase neurogenesis during development.

### Nervous system-specific induction of YFs leads to cortical expansion during development

To remove any detrimental effects of ubiquitous YFs induction during development, i4F mice were crossed with a Cre-dependent floxed rtTA and the nervous system-specific Nestin-Cre mouse line. This mouse model (i4F-Nes) allows the study of high and longer Dox treatments during development and at later time points (Fig. 2A). We found that high Dox concentrations, or longer treatments, resulted in much bigger brains, with an expanded cortex and weight of up to three times the size of their control littermates (CTR) at E17.5 (Fig. 2B-C, S2A). Moreover, we observed strong induction of YFs such as c-Myc, Oct4 and Sox2, together with Nanog expression in both germinal and neuronal layers after high Dox treatment (Fig. S2B-C). Upon further inspection of i4F-Nes embryos treated with different conditions, we focused on the low Dox treatment (4d, E10.5-E14.5) because it showed complete preservation of brain morphology and embryo survival to adult stages, as in the i4F-Rosa mouse line. We quantified the numbers of apical (Pax6+) progenitors and also stained for Pvim and PH3. We found increased numbers of basal PH3+ and Pvim+ cells in i4F-Nes embryos at E15.5 (Fig. 2E). Moreover, the total numbers of Pax6+ progenitors were increased (Fig. S2D). This change was mainly due to the presence of Pax6+ cells in neuron-enriched layers (IZ+CP) (Fig. 2E). These results, together with an increased perimeter of lateral ventricle walls and reduced thickness of the ventricular zone (Fig. 2E), were similar to those found in the i4F-Rosa line with the same treatment (Fig. 1I-J). This suggests that the observed changes in i4F-Rosa brains can be largely explained by the effects of YFs induction in the nervous system.

**Figure 2.**
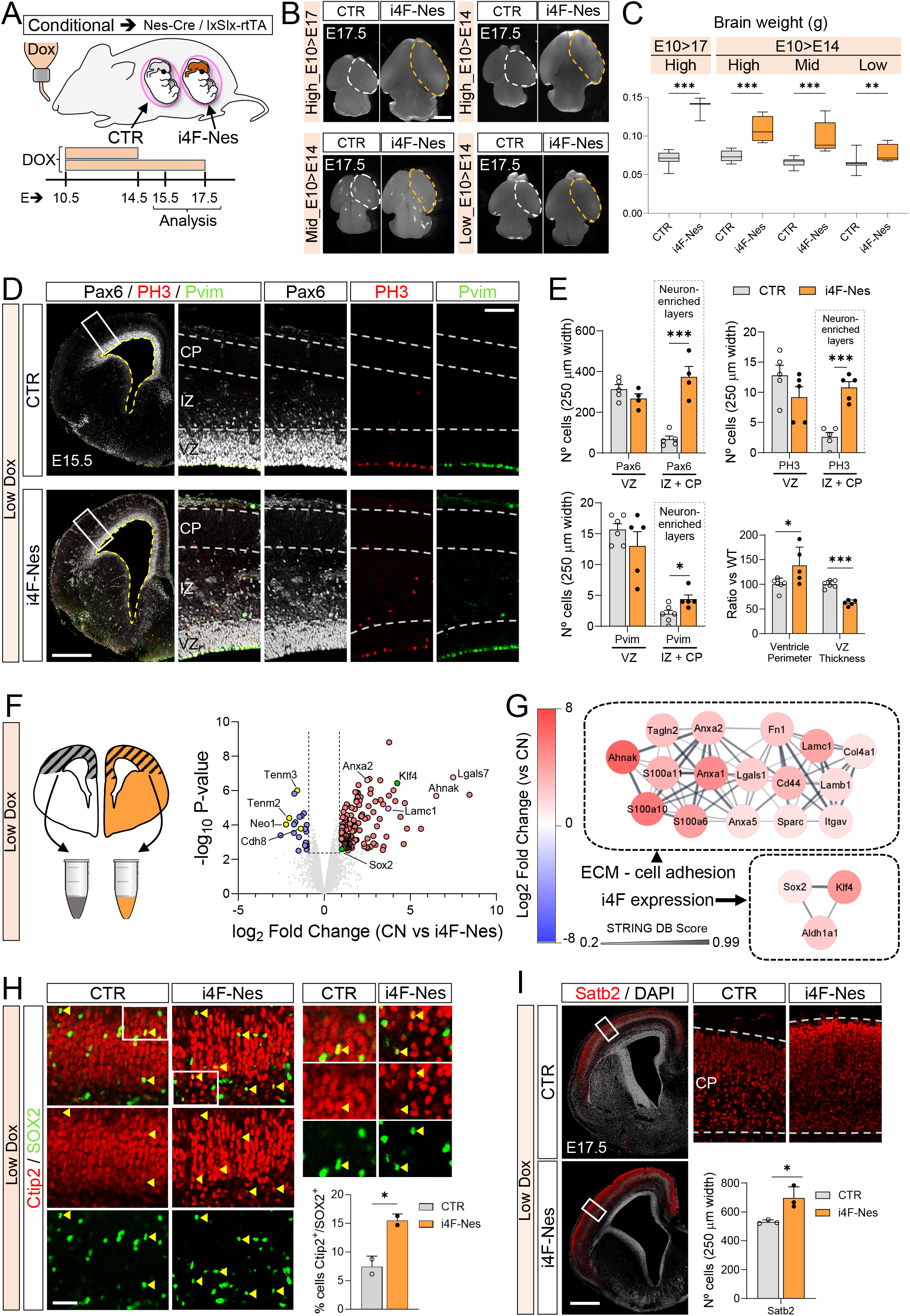
Nervous-system specific induction of YFs leads to cortical expansion during development. (A) Scheme of Dox treatment in control (CTR) and i4F-Nes littermate embryos used in downstream analysis. (B) Top view of E17.5 mouse brains from control (CTR) and i4F-Nes embryos with different Dox treatments. Dashed lines delineate the cortical area of one hemisphere. (C) Brain weight of data shown in (B). n=5-6 mice/group. **p<0.01,***p>0.001, unpaired Student’s t test. (D) E15.5 coronal brain sections, after 4d of Low Dox (E10.5 to E14.5), were labeled for neuronal progenitors (Pax6, white), dividing RG cells (Pvim, green) and a mitotic cell marker (PH3, red). Areas in dashed rectangles are shown with higher magnification on the right. (E) Quantification of data shown in (D). n=4-6 mice/group. *p<0.05,***p<0.001, unpaired Student’s t test. (F) Scheme showing the E15.5 cortical area used for mass spectrometry experiments and Volcano plot showing down-regulated (left, blue) and up-regulated (right, red) proteins in i4F-Nes cortices respect to CTR samples. Surface receptors are colored in yellow and YFs in green.(G) Cytospace Network model showing protein-protein interactions (PPIs) of 19 up-regulated proteins in i4F-Nes samples from data shown in (F). The color bar represents the Log2 Fold change protein ratios. The edges represent PPIs obtained from the STRING database (H) Similar experiment as shown in panel D, except for cortical plate insets stained with the neuronal marker Ctip2 (red), and progenitor marker Sox2 (green). White rectangles are magnified on the right. Yellow arrowheads indicate double positive Ctip2/Sox2 cells, and their quantification is shown on the graph. n=2 mice/group. *p<0.05, unpaired Student’s t test. (I) Similar experiment as shown in panel D, except for E17.5 coronal brain sections stained with upper cortical neuronal marker Satb2 (red) and DAPI (White). White rectangles are magnified on the right. The graph shows the quantification of the number of Satb2+ cells in the cortical plate. n=3 embryos/group. *p<0.05, unpaired Student’s t test. The data are represented as mean ± SEM. Circles indicate values of individual brains. Scale bars, 1mm (B), 500um (D,I), 100um (H).

A recent study has linked increased adhesion in proliferative neural stem cell niches to reduced proliferative potential, a process that naturally occurs during aging^29^. We performed a proteomic profiling of cortices from E15.5 i4F-Nes and their corresponding control littermates (CTR) after 4d of low Dox treatment (Fig 2F). As expected, we found significantly increased protein levels of YFs such as Sox2 and Klf4. Interestingly, we also found reduced expression of cell adhesion proteins such as Cdh8 and Neo1, and changes in extracellular matrix components (ECM) in i4F-Nes cortices (Fig. 2F-G). These data suggest that YFs induction alters the properties of the neuronal progenitor niche, making it less adhesive and thereby favoring proliferation during development, in addition to influencing cell identity. Indeed, similar to Pax6, we found ectopic expression for Sox2 in neuronal-enriched layers, where several Sox2+ cells co-expressed neuronal markers such as Ctip2 (Fig. 2H). The expansion of neuronal progenitors resulted in an increased number of upper cortical neurons at E17.5 (Fig. 2I). Taken together, these results indicate that conditional nervous-system YFs induction transiently perturbs cell behavior leading to expansion of progenitors and consequently to increased output of upper cortical neurons.

### Improved behavioral performance after transient YFs induction during development

The increases in cortical thickness and cortical neuron numbers observed in embryos from both i4F-Rosa and i4F-Nes mouse lines were found to persist into adulthood (Fig.3 B-C). The increase in CP thickness during development was due to specific thickening of upper cortical layers. Consistent with this, adult (5-month-old) i4F-Rosa mice showed increased numbers of neurons and an enlarged layer II-IV (Fig. 3B-C). Because of the enlarged cortex tissue, the increased numbers of upper layer neurons were not accompanied by alterations in cell density. We next wondered about the functional consequences of these alterations and conducted an array of behavioral tests at adult stages (Fig. 3A).

**Figure 3.**
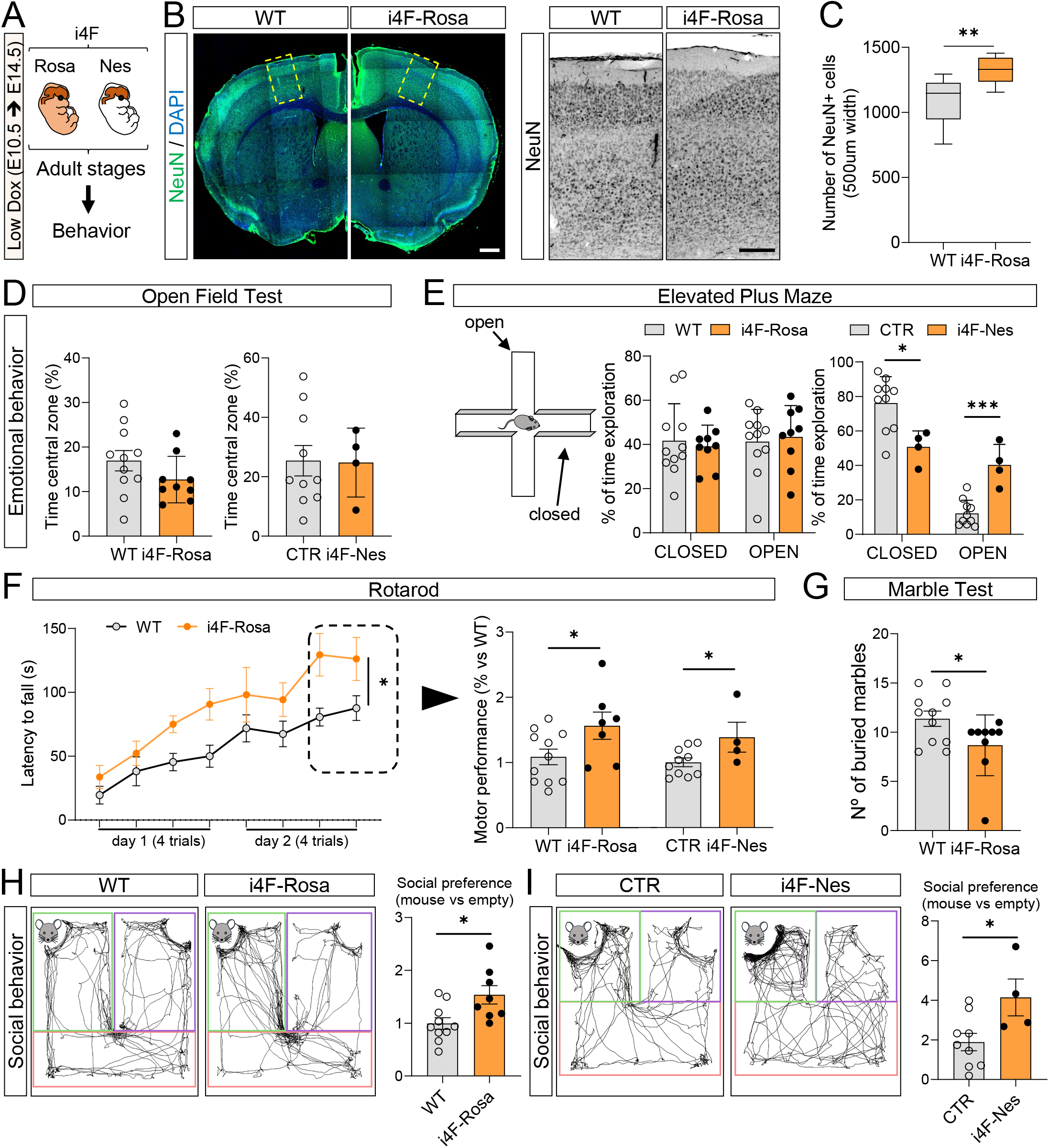
Increased behavior performance after transient YFs induction during development. (A) Scheme of mice used for behavioral experiments at adult stages (4-5 months old). i4F-Rosa and i4F-Nes mutant embryos and their corresponding controls (WT and CTR) were treated for 4d with Low Dox during development. (B) Adult coronal sections stained with the neuronal marker NeuN (green) and DAPI (nuclei, blue). Areas in dashed rectangles are shown with higher magnification on the right. (C) Quantification and whisker plot of the data shown in (B). n=9-11 mice/group. **p<0.01, unpaired Student’s t test. (D) Percentage of time spent at the central zone of i4F-Rosa (9), i4F-Nes (4) and their respective controls WT (11) and CTR (10). No significant changes between groups, unpaired Student’s t test. (E) Percentage of time spent in closed and open arms in the elevated Plus Maze test. n=11 (WT), 9 (i4F-Rosa), 10 (CTR) and 4 (i4F-Nes). *p<0.05, ***p<0.001, Two-way ANOVA. (F) Quantification of the latency to fall in the accelerating rotarod task (4 sessions per day during 2d).n=11 (WT) and 9 (i4F-Rosa). *p<0.05, Two-way ANOVA. The average of the last two sessions per mice (both i4F-Rosa and i4F-Nes with their controls) is shown on the right. n=11 (WT), 9 (i4F-Rosa), 10 (CTR) and 4 (i4F-Nes). *p<0.05, unpaired Student’s t test. (G) Number of buried marbles during 20min session in the Marble burying test. n=11 (WT), 9 (i4F-Rosa), *p<0.05, unpaired Student’s t test. (H) Social preference index in the three-chamber social interaction test. Representative mice track paths and the location of the target mouse (green area), empty cage (magenta area) and connecting chamber (red area). Ratios are calculated based on the time spent on the target mouse vs empty cage. n=11 (WT), 9 (i4F-Rosa), *p<0.05, unpaired Student’s t test. (I) Similar experiment as shown in panel H, except for i4F-Nes and their controls (CTR). n= 10 (CTR) and 4 (i4F-Nes). *p<0.05, unpaired Student’s t test. The data are represented as mean ± SEM. Circles indicate values of individual mice. Scale bars, 500um (B), 200um inset (B).

Higher cognitive abilities are thought to arise from cortical expansion during evolution^30,31^. We thus tested whether cortical expansion by YFs induction could affect specific behavioral parameters. We subjected 5-month-old i4F-Rosa and i4F-Nes mice, with their respective littermate controls (WT and CTR, respectively), to a battery of behavioral tests targeting different features. The data obtained from the open field test showed no significant differences of the exploration habits between i4F-Rosa and i4F-Nes and their controls (Fig. 3D). Next, we performed the elevated-plus maze test to determine whether mutant mice showed altered anxiety-like behavior. There were no significant differences in the times spent in the open and closed arms between i4F-Rosa and WT mice. Notably, i4F-Nes mice showed a significantly increased time exploring the open arms (Fig. 3E). These results suggest that transient induction of YFs during development has no effect (i4F-Rosa) or even reduces (i4F-Nes) anxiety-like behavior at adult stages (Fig. 3E). We next evaluated higher-order cognitive functions that are related to cortical function such as motor learning, compulsive-like behavior and sociability^32,33^. We first performed the accelerating rotarod test to evaluate motor learning^34^. Importantly, i4F-Rosa mice showed faster and an enhanced ability to maintain their balance compared to WT mice (Fig. 3F). The average of the last two sessions, during which a plateau of learning was reached, was used to compare their motor performance. Both i4F-Rosa and i4F-Nes mice exhibited improved motor performance with respect to their controls (WT and CTR, respectively) (Fig. 3F). Then, we performed the marble burying test to evaluate potential compulsive-like behaviors^35^. i4F-Rosa mice showed a reduced number of completely buried marbles compared with their WT littermates (Fig. 3G), indicating reduced compulsive behavior. Finally, to evaluate sociability, we performed the three-chamber social interaction test^36^. In this paradigm, both i4F-Rosa and i4F-Nes mice showed an increased sociability index when compared with their respective controls (WT and CTR) (Fig. 3H-I). We conclude that cortex expansion by transient YFs induction during development improves motor and social behavior of adult mice.

### Intermittent YF induction in adult hippocampal principal neurons is tolerated and prevents synaptic loss in the 5xFAD mouse model

YFs have been shown to reverse some aging features and increase regeneration in mammalian tissues^2,37–39^. Since our data showed that transient YFs induction is tolerated by post-mitotic neurons during development, together with enhanced cognitive skills, we asked whether applying a similar induction protocol at adult stages could be beneficial in protecting against neurodegeneration. We chose the 5xFAD mouse model, a robust model of Alzheimer disease (AD), with neurodegeneration and cognitive deficits^40^. We established an *in vivo* intermittent YF induction protocol using similar Dox treatment that was tolerated by neurons during development. 5xFAD mice were crossed with the i4F mouse line generating the i4F/5xFAD mouse line and its littermate controls (i4F). Notably, in this mouse line, YFs induction was driven by Synapsin-1 (SYN1)-dependent rtTA-expressing adeno-associated virus (AAV) injected into the dorsal hippocampus of eight-week-old mice (Fig. 4A). We introduced a ZsGreen reporter driven by a tetracycline operator (TRE) promoter to visualize neurons expressing YFs after Dox treatment. From 12 to 35 weeks, both groups, (i4F/5xFAD and its control i4F), were treated with vehicle (VEH) or low Dox (0.2mg/ml in the drinking water) 3d ON, 4d OFF per week (Fig. 4A). After treatment (8-month old), all mouse groups were subjected to comprehensive behavioral, histological and molecular experiments (Fig. 4A).

**Figure 4.**
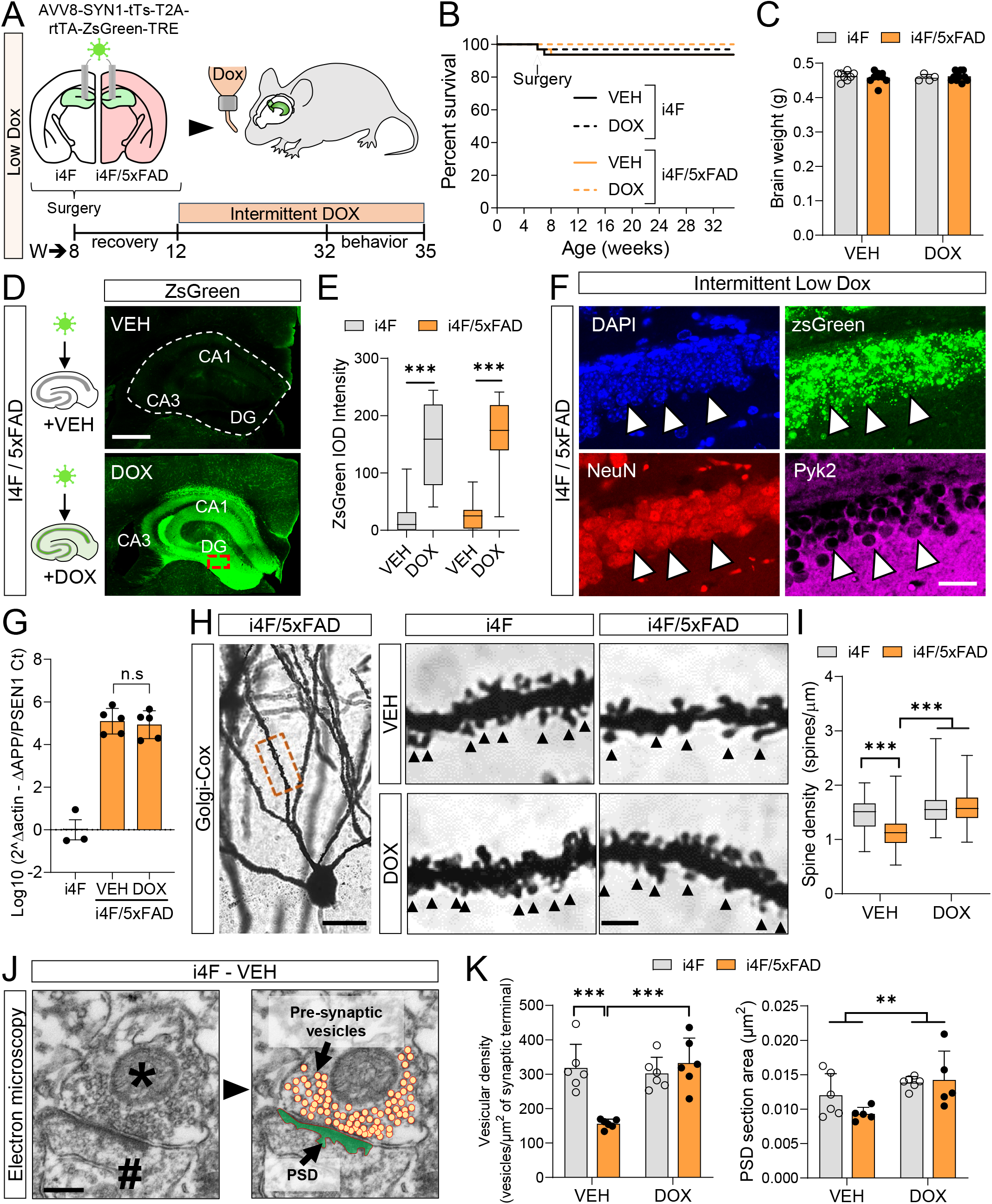
Establishment of in vivo YF induction protocol prevents synaptic loss in the 5xFAD mouse model. (A) Experimental design schematic and delivery of AAV-SYN1-tTs-T2A-rtTA-ZsGreen-TRE into the hippocampus of i4F and i4F/5xFAD mice. Both groups were treated with intermittent Dox or vehicle (VEH) for 5 months. (B) Survival outcome for each group during experimental protocol (n=15-20 mice/group). Only three deaths occurred due to post-surgical complications. No significant changes between groups, Kaplan-Meier method. (C) Quantification of total brain weight at the end of the experimental protocol. n=15-20 mice/group. No significant changes between groups, Two-way ANOVA with Bonferroni’s post hoc test. (D) Validation of rtTA-dependent ZsGreen reporter signal after 5 months Dox and VEH intermittent treatment. Intensity of Zsgreen signal (green) in coronal hippocampal sections of Dox and VEH treated i4F/5xFAD mice (8-month-old). Red dashed rectangle is shown with higher magnification on panel F. (E) Whisker plot of Zsgreen intensity in each group. n=15-20 mice/group. ***p<0.001 DOX vs VEH in each group, Two-way ANOVA with Bonferroni’s post hoc test. (F) Inset from panel D showing Zsgreen signal (green) was stained with NeuN (mature neuron, red), Pyk2 (mature principal neuron, magenta) and DAPI (nuclei, blue). (G) qRT-PCR for APP/PSEN1 transgene in the hippocampus of i4F and i4F/5xFAD treated with VEH or Dox for 5 months. Expression normalized to housekeeping gene Actin and presented as whisker plots from 3-5 mice/group (H) Golgi-Cox-stained granular neuron located in the dentate gyrus molecular layer. Red dashed rectangle indicates the region of the secondary apical dendrite that was analyzed. This region is shown with higher magnification on the right. Black arrowheads indicate dendritic spines. (I) Quantification of data shown in G. n=7-11 mice/group (up to 100-150 dendrites per group). ***p<0.001, Two-way ANOVA with Bonferroni’s post hoc test. The data are presented as whisker plots. (J) Electron microscope image of an excitatory synapse in the molecular layer of the dentate gyrus in 8-month-old i4F-VEH. Asterisk indicates the presynaptic component; hashtag indicates the postsynaptic component. Presynaptic vesicles are coloured with orange circles; post-synaptic density (PSD) is coloured in green. (K) Quantification of the vesicular density and PSD area in all groups. n=6 mice/group. **p<0.01, ***p<0.001, Two-way ANOVA with Bonferroni’s post hoc test. The data are represented as mean ± SEM. Circles indicate values of individual mice. Scale bars, 400um (D), 100um (F), 15um, 3um (inset) (H) and 0.25um (J).

Our intermittent YFs induction was well tolerated, and survival was not affected in any experimental group (Fig. 4B). No treatment-dependent changes in body or brain weight were found at 8 months (Fig. 4C). We then assessed the gross hippocampal anatomy as well as the neuronal identity of the targeted cells. First, our intermittent YFs induction did not affect the hippocampal formation in terms of organization and gross anatomy (Fig. 4D). Targeted neurons with correct activation of the TRE promoter were found expressing ZsGreen in the Dox-treated groups (Fig. 4D-E). Second, immunostaining for two markers of mature neurons, NeuN and Pyk2, showed that the identity of the transduced granule cells of the dentate gyrus (DG) was not altered by YFs induction (Fig. 4F). Third, quantitative RT-PCR (qRT-PCR) analysis confirmed that intermittent reprogramming does not alter *per se* the expression of the APP/PSEN1 transgene present in the 5xFAD mouse model at 8 months(Fig. 4G).

We then evaluated key aspects of structural plasticity in granular neurons by Golgi staining. We found the expected reduction in the spine density of secondary dendrites in i4F/5xFAD treated with VEH (i4F/5xFAD-VEH) compared with the control i4F-VEH group (Fig. 4H). Notably, this reduction was completely rescued in the i4F/5xFAD group treated with Dox (i4F/5xFAD-Dox) (Fig. 4I). Next, we further examined presynaptic and postsynaptic changes using electron microscopy. We quantified the numbers of presynaptic vesicles per synapse and the post-synaptic density (PSD) area in the dentate gyrus (DG) molecular layer of the four groups of mice at 8 months of age (Fig. 4J). The numbers of presynaptic vesicles per synapse were significantly decreased in the i4F/5xFAD-VEH, as described previously^41^, but they were completely rescued in the i4F/5xFAD Dox group (Fig. 4K). We additionally found that the PSD area was significantly increased in both mouse groups (i4F and I4F/5xFAD) subjected to intermittent reprogramming (Fig. 4K). In summary, these results revealed that intermittent YF induction is tolerated by adult hippocampal neurons and induced synaptic improvements in the 5xFAD mouse model.

### Amelioration of AD-related hippocampal plaques and proteomic signatures by YF induction

Because 5xFAD mice develop severe amyloid pathology and proteomic changes in the hippocampus from 2 to 8-months of age^40,42^, we investigated whether intermittent YFs induction during this period may have beneficial effects on these hallmarks. We quantified the plaque load in the three main dorsal hippocampal subfields, namely CA1, CA3 and DG (Fig. 5A), by Aβ immunolabelling. The number (Fig. 5B) and size (Fig. S3) of plaques in these regions were strongly reduced in i4F/5xFAD Dox with respect to i4F/5xFAD-VEH mice . The plaque numbers were not altered in other regions without YFs induction, such as the cortex, in the i4F/5xFAD-Dox mice (Fig. 5B). To identify proteomic differences between the groups, we performed quantitative proteomic profiling of the DG. This region is critical for learning and memory, and it is one of the most affected hippocampal areas in AD^43^. Proteomic analysis showed that the transgene expression of *APP* present in the 5xFAD mouse was not affected by YFs induction at 8 months (Fig. 5D), which was consistent with qRT-PCR experiments (Fig. 4G). Other AD-related markers such as neuroinflammation and stress responses were all unchanged in adult i4F/5xFAD-Dox compared with i4F/5xFAD-VEH group (Fig. 5D). Next, we identified differentially expressed proteins (DEPs) between i4F/5xFAD-VEH and their littermate controls i4F-VEH. We found 950 DEPs in i4F/5xFAD-VEH with respect to its i4F-VEH control. The 20 most enriched KEEG pathways were predicted using Fold Enrichment analysis with the ShinyGO tool^44^ and visualized as an interaction network using Cytospace^45^ (Fig. 5E). The significant pathways found in the DG of i4F/5xFAD-VEH were functionally related to neurodegenerative processes (Fig. 5E), as described previously^42^. Fold change enrichment analysis showed that 19 out of the 20 most significantly enriched pathways altered in i4F/5xFAD-VEH vs i4F-VEH were partially rescued in the i4F/5xFAD-Dox group, suggesting that intermittent YF induction ameliorates proteomic changes present in the 5xFAD mouse model (Fig. 5F). Among 950 DEPs found in the i4F/5xFAD-VEH, 493 protein levels were normalized in the i4F/5xFAD-Dox compared with i4F-VEH control group. To explore the most relevant collective functions of the rescued DEPs in the i4F/5xFAD-VEH, we constructed a network model to describe their interactions. We focused on 103 DEPs whose network has been found to be related to AD^42^. The results showed that the rescued pathways in i4F/5xFAD-Dox group were related to the pathogenesis of AD (mitochondria function, cellular metabolic processes, cellular adhesion, degradation and coagulation) (Fig. 5G). Together, these findings suggest that intermittent YFs induction ameliorates hippocampal plaque loading and rescues several proteomic signatures present in the 5xFAD mouse model.

**Figure 5.**
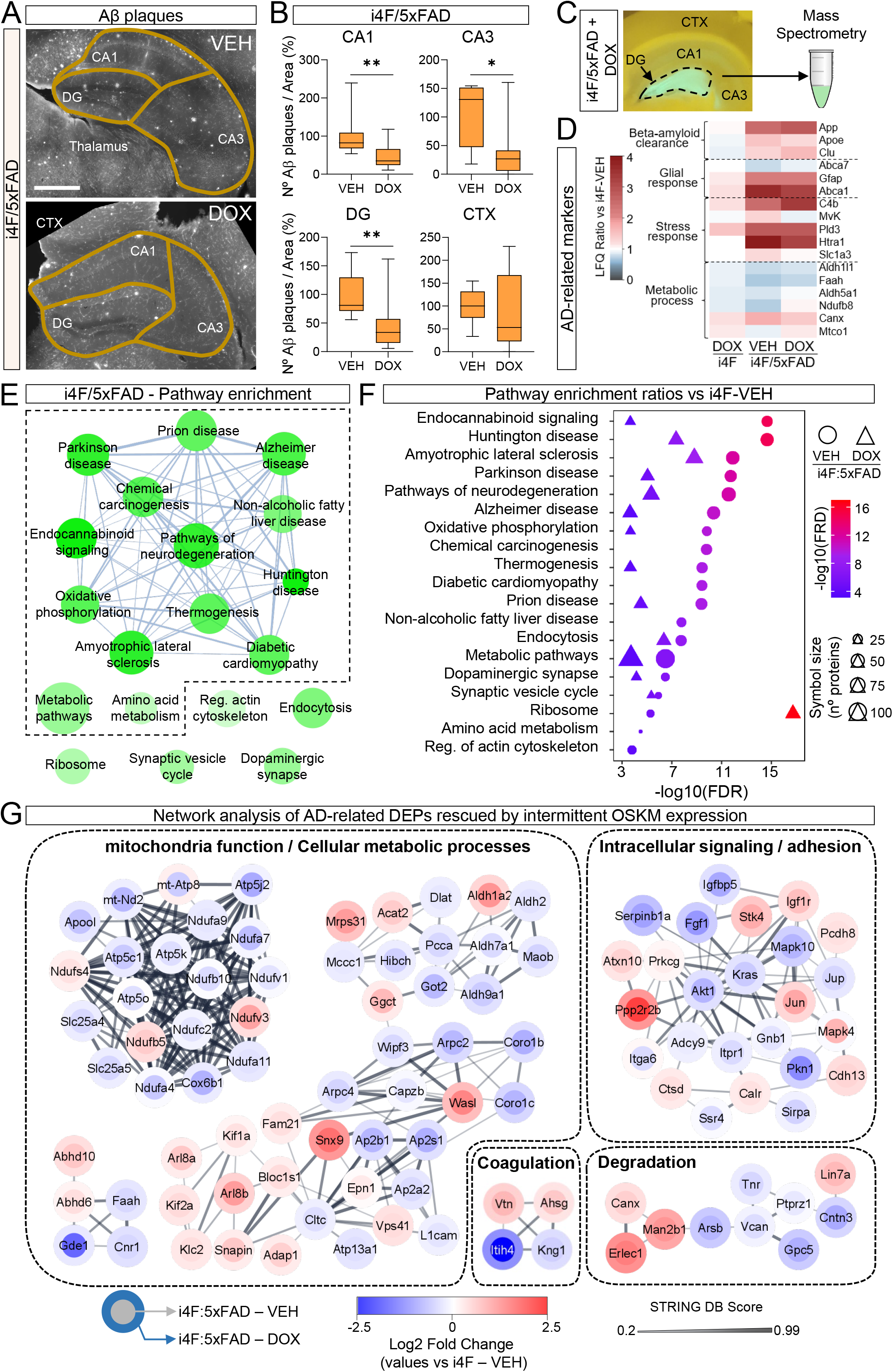
Amelioration of AD-related hippocampal plaques loading and proteomic signatures by YF induction. (A) Coronal hippocampal sections of i4f/5xFAD mice(-VEH and -DOX) were labeled for Aβ immunoreactivity. Hippocampal CA1, CA3, dentate gyrus (DG), and motor cortex sub-regions were delineated following the Gaidi mouse brain atlas. (B) Quantification of number of plaques in each region as shown in (A). n=5-7mice/group. Three slices per mouse (separated by 240um, from -1.28 to -2mm from Bregma). *p<0.05, **p<0.01, unpaired Student’s t test. (C) Scheme showing the location of the DG used for proteomic profiling experiments. (D) Relative protein expression levels across different groups respect to control (i4F-VEH). Protein expression levels based on untargeted label-free quantitation (LFQ) was normalized to i4F-VEH control group (value =1). The color bar represents the gradient of normalized protein abundances. (E) Network analysis of enriched KEEG pathways of differential expressed proteins (DEPs) of i4F/5xFAD-VEH respect to i4F-VEH group. Significant enriched pathways (Fold enrichment, FDR, p<0.05) are shown. n=4-6 mice/group. (F) Fold enrichment analysis of KEEGs pathways shown in E for VEH and DOX treated i4F/5xFAD mice respect to i4F-VEH group. n=4-6 mice/group. (G) Cytospace Network model showing protein-protein interactions (PPIs) of 103 DEPs of i4F/5xFAD-VEH respect to i4F-VEH group that are rescued in the i4F/5xFAD-DOX group. The color bar represents the Log2 Fold change protein ratios. Node color represents an increase (red) or decrease (blue) in i4F/5xFAD DOX (center) and VEH (boundary) compared to i4F-VEH (control group). The edges represent PPIs obtained from the STRING database. Scale bar, 450um (A).

### YF induction prevents cognitive decline in the 5xFAD mouse model

The recovery of several AD-related hallmarks of the 5xFAD mouse model with intermittent YFs induction (i4F/5xFAD-Dox group), prompted us to explore whether there was an improvement in the cognitive deficits present at 8 months of age in hippocampal-related tasks^40,46–48^. We conducted different behavioral tests, including emotional behaviors (plus maze or PM and forced swimming test or FST), cognitive flexibility and spatial memory (T-maze), aversive associative memory (Passive avoidance), spatial long-term (NOLT) and working (Y-maze) memories (Fig. 6A). The data obtained from the plus maze (Fig. 6B) and forced swimming test (Fig. 6C) revealed statistically significant differences between i4F/5xFAD (VEH and Dox groups) and their respective controls (i4F-VEH and -Dox), but no changes between i4F/5xFAD treated with Dox or VEH. These results indicate that although the 5xFAD genotype is associated with disturbances on anxiety or mood-related behaviours, they were not modified by YF induction in the dorsal hippocampus. This result agrees with studies linking emotional-like behaviors rather to the ventral hippocampus, whereas the dorsal region is principally involved with cognitive functions^49^.

**Figure 6.**
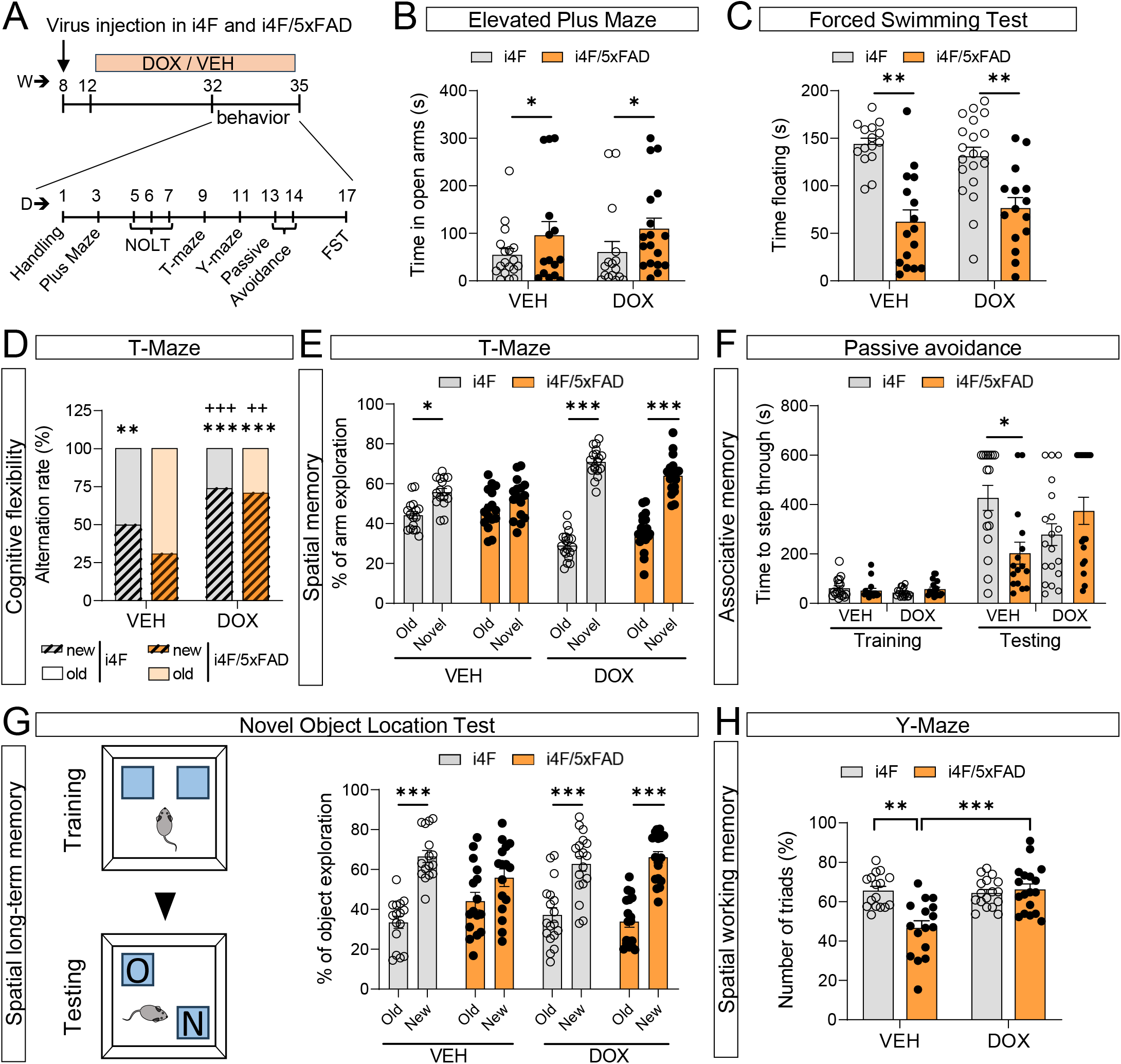
YF induction prevents the cognitive decline in the 5xFAD mouse model. (A) Experimental design and sequence of behavioral tests performed. (B) Percentage of time spent in open arms in the elevated Plus Maze test. n=15-20 mice/group. *p<0.05 between genotypes, Two-way ANOVA. (C) Time floating in the forced swimming test. n=15-20 mice/group. **p<0.01 between genotypes, Two-way ANOVA. (D-E) Quantification of cognitive flexibility and Spatial memory in a T-maze. (D) Cognitive flexibility was measured as the first arm visited in the testing trial. Alternation is presented as %. n=15-20 mice/group. **P < 0.01, ***P < 0.001 respect to i4F/5xFAD-VEH. ++P < 0.01,+++P < 0.001 respect to i4F-VEH. Chi-square test. (E) Percentage of time exploring the novel arm (new context) versus the time exploring the old arm (old context). n=15-20 mice/group. *P < 0.05, ***P < 0.001 comparing % of novel arm vs old arm preference. Two-way ANOVA with Bonferroni’s post hoc test. (F) Associative learning measured using the passive avoidance paradigm. Quantification of the latency (seconds) to step-through in both training and testing trial for all groups after receiving an electric shock (2 s/1 mA). n=15-20 mice/group. *p<0.05 respect to i4F-VEH group. Two-way ANOVA with Bonferroni’s post hoc test. (G) Spatial long-term memory using the novel object location test. Percentage of time exploring the object placed in a new location (New) versus the time exploring the object placed in an old location (Old). n=15-20 mice/group. ***p<0.001 comparing time exploring objects in new vs old location. Two-way ANOVA with Bonferroni’s post hoc test. (H) Quantification of spontaneous alternation in the Y-maze. n=15-20mice/group. **p<0.01, ***P < 0.001 respect to i4F/5xFAD-VEH group. The data are represented as mean ± SEM. Circles indicate values of individual mice.

Next, we applied different tasks to compare memory performance in these mice. We first evaluated cognitive flexibility and spatial memory in a T-maze (Fig. 6D-E). We found that i4F/5xFAD-VEH mice alternated significantly less than i4F-VEH control group, therefore displaying cognitive inflexibility. Notably, both the i4F-Dox and i4F/5xFAD-Dox groups with intermittent YFs induction, performed better, i.e. more flexibly, than i4F-VEH group (Fig. 6D). We also quantified the time a mouse spends exploring a familiar vs a novel arm. i4F/5xFAD-VEH mice displayed no preference for the novel arm. This effect was completely rescued in the i4F/5xFAD-Dox group (Fig. 6E). We then examined associative memory in the passive avoidance task. Latency to step-through during the training session was similar between groups (Fig. 6F). However, in the testing session, although all groups showed a significant increase in the latency to enter the dark compartment 24 h after receiving an electrical shock, this latency was shorter in i4F/5xFAD-VEH mice compared with i4F-VEH control group, and completely rescued in i4F/5xFAD-Dox mice (Fig. 6F). The novel object location test also showed that i4F/5xFAD-Dox had rescued the preference for novel objects that is lost in the i4F/5xFAD-VEH mice (Fig. 6G). Finally, mice were tested in the spontaneous alternation in a Y-maze paradigm that assesses spatial working memory^50^. The i4F-VEH and i4F-DOX mice displayed a spontaneous alternation over the chance level (∼65%), whereas the arm choice was decreased to ∼50% (chance levels) in the i4F/5xFAD-VEH mice (Fig. 6H). The spontaneous alternation was restored in the i4F/5xFAD-Dox mice (Fig. 6H). Additional controls testing the effects of Dox treatment alone showed no differences in either WT or 5xFAD mice (Fig. S4). In summary, YF induction in principal neurons of the dorsal hippocampus in the 5xFAD mice prevented their cognitive decline but not their emotional alterations.

## Discussion

In this study, we demonstrate that transient reprogramming with YFs not only safely increases neurogenesis during mouse brain development but also prevents the development of AD-related features in adulthood. The enhancement of neurogenesis during development results in a greater production of neurons and expansion of the cortex, both of which correlate with improved behavioral performance. At adult stages, we found that principal neurons in the hippocampus tolerate transient YFs expression for several months without losing their cell identity. Instead, the expression of YFs prevented the development of several AD-related hallmarks in the 5xFAD mouse model and protected against cognitive decline. These findings enhance our understanding of YFs as a tool to modulate neurogenesis and highlight their potential use in brain disorders.

### Increased neurogenesis and cortex expansion by transient YFs expression

We present evidence that YFs can perturb cell identity of both progenitors and neurons during brain development. Strong induction of these factors leads to a reduction in Pax6 expression in apical progenitors. It also results in a massive ectopic expression of both Pax6 and Sox2 in neuronal layers, where their expression is low in control conditions. Moreover, long-term induction of YFs during development can affect cell fate. This is evidenced by the loss of identity markers such as Pax6 for progenitors and Ctip2 for neurons, along with the emergence of Nanog expression, a well-known pluripotent marker. This extends the concept that strong and continued overexpression of these factors can induce pluripotency in various cells from peripheral tissues *in vivo*^3,17^.

Importantly, both progenitors and neurons tolerate low and transient YFs expression, leading to increased neurogenesis and a higher number of upper cortical neurons without the formation of tumors. Their identity is temporarily perturbed by the coexpression of these factors, but it is not lost. This observation aligns with the notion that transient de-differentiation is proposed as an integral step in the reprogramming process^51,52^. In line with this model, cells undergo transient de-differentiation post YFs induction, and once their expression stops, revert to their original cellular identity. Recent studies, which employ transient reprogramming protocols to rejuvenate multiple murine cell types, further confirm this. In these studies, cells display a temporary identity perturbation when YFs are expressed. However, this identity is recovered, likely due to epigenetic memory or persistent expression of specific identity genes^53,54^. The increased neurogenesis goes in line with previous studies showing increased proliferation after YFs induction in peripheral tissues^11^

In the cortex, we identified widespread proteomic changes associated with ECM following transient YF expression during development. Previous studies have shown that their induction alters the expression of key components involved in cell adhesion, cell cycle, protein trafficking and extracellular matrix^55,56^. Interestingly, some of these alterations, including the reduced expression of proteins involved in cell adhesion such as Cdh8 and Neo1, could be related with the expansion in neurogenesis that we observed during development. Indeed, Cdh8 is expressed in the germinal zone during development, and its knockdown amplifies both proliferation and neuron numbers^57^. Similarly, genetic ablation of Neo1 also increases proliferation^58^. Moreover, a recent study indicated that aging has a profound effect on the expression of genes involved in cell-cell adhesion and cell-matrix interactions in brain stem cell niches. Aging increases the cellular adhesion of proliferative neural stem cells, which correlates with reduced neurogenesis potential^29^. Accordingly, our results suggest that transient reprograming could also rejuvenate the niche of neuronal progenitors by reducing the expression of cell adhesion proteins, thereby increasing their neurogenesis capability.

The finding that transient reprogramming during development leads to an increase of upper-layer neurons and cortical expansion that persist into adulthood, raised the question to what extent this could improve behavioral performance in adult mice. Most studies that analyze the effects of genes associated with cortical expansion in mouse embryos have not examined the impact on cognitive abilities at adult stages^59^. Two recent reports achieving around 1.2-fold increase in upper-layer neurons in adult mice, have found improved hippocampus-independent memory flexibility^60,61^. We were surprised to find that our mouse model displays the strongest increase in the number of upper-cortical neurons (up to 1.4-fold) at adult stages ever reported, particularly in the motor and prefrontal cortical areas. This is consistent with the massive expansion of rostral cortical areas during development (Fig. 2B and S2A), and with improved behavior in related tasks at adult stages. For example, we observed enhanced performance on the accelerated rotarod, a well-established paradigm for cortex-dependent motor skill learning^62^. This is fully in line with the increased abundance of upper-layer neurons in the motor cortex, which is known for its critical role in motor skill acquisition^63^. Similarly, we noted improved social behavior, predominantly associated (but not limited to) with the prefrontal cortical region^64^, which shows expansion following transient reprogramming. Interestingly, the human prefrontal cortex is the most expanded cortical region^65^. Several studies provide strong evidence that such expansion and improved social cognition is a key step in the evolution of human intelligence^66^.

### Transient YF reprogramming is protective in an Alzheimer Disease mouse model

Previous studies have demonstrated beneficial effects of intermittent YFs expression in various contexts in adult mice related to regeneration and aging^2,5,37,67^. In vivo partial reprogramming alters age-associated molecular changes during physiological aging in mice^2^. However, in contrast to the present work, none of these studies has explored its impact on mature principal neurons in the context of neurodegeneration. Our study presents a novel application that prevents the development of several neurodegenerative hallmarks as well as cognitive decline in a well-established mouse model of AD. Importantly, these improvements were observed without any discernible impact on the general health of the mice or the identity of targeted principal neurons in the hippocampus. This aligns with prior seminal studies highlighting the safety of transient reprogramming^2,4^.

Dendritic spine pathology and the loss of synaptic contacts are defining features of neurodegenerative disorders, including AD^68–71^. We found that intermittent YFs expression rescued several synaptic features, such as spine density and pre-synaptic vesicular density, at least in granular hippocampal neurons. This result is consistent with a recent observation that YFs induction increases the level of the synaptic protein GluN2B, particularly in dendrites, in the dentate gyrus ^26^. This protein is known to facilitate synaptic potentiation and to promote dendritic spine density and function^72^ . Moreover, we observed reduced Aβ loading after YFs induction in the 5xFAD mouse model. Previous studies have shown that these factors can affect the expression of several promoters, including Thy1^54^. Our qRT-PCR and proteomic analysis show that the effects we observed, such as in Aβ loading, are not due to altered expression of the APP/PSEN1 transgene, or changes in AD-related glia response (Fig.5D). Furthermore, our proteomic network analysis identified mitochondria function and metabolism as one of the most relevant modules affected in the 5xFAD mouse model. This result is consistent with a recent transcriptomic study from human AD patients^73^, and previous proteomic analysis from AD mouse models^74^ and human patients^75^. Another significant pathway that was rescued is the one related to the endocannabinoid system, which agrees with previous studies highlighting the potential of this pathway in the modulation of Aβ loading and synaptic dysfunction^76^. Interestingly, several of these proteomic AD-related signatures were rescued by transient YF reprogramming, including proteins that are strongly associated with AD in humans.

Aging critically influences both Aβ loading and mitochondrial function. For example, neurons are the primary source of Aβ, and a recent report indicates that its production and accumulation increase with age^77^. Blocking Aβ production in aged neurons prevented the decline in their synapses^77^. Regarding mitochondrial function, aging correlates with a decline in its function, as shown by reduced ATP production, lipid metabolism and biogenesis^78^ . Given our findings that transient reprogramming improves synaptic plasticity in hippocampal neurons, combined with amelioration of Aβ loading and mitochondrial proteomic signatures in the 5xFAD mouse model, suggest that this process may rejuvenate these neurons. This is consistent with the prevailing hypothesis that partial reprogramming can rejuvenate cells^79,80^. Our results could be an interesting starting point for further research.

This study also revealed that partial reprogramming prevents cognitive decline in the 5xFAD mouse model. This includes behavioral paradigms related to associative learning, cognitive flexibility, spatial memory, and spatial working memory previously described to be affected in the 5xFAD mice^81–83^. These improvements are associated with the YFs expression in hippocampal granular and pyramidal neurons. Indeed, our proteomic analysis and histology did not identify significant changes in other cell populations, such as microglia and astrocytes (data not shown). Although YFs induction in mature neurons might alter several mRNA transcription processes, as demonstrated in other systems^56^, it is conceivable that the rescue in spine density, Aβ loading, and mitochondrial proteomic features contribute to better cognitive performance in the 5xFAD mouse model. It is interesting to find that in some behavioral paradigms, such as cognitive flexibility, both control and 5xFAD mouse subjected to partial reprogramming outperformed their respective controls. Future studies should investigate the specificity or generality of YFs induction impact on neuronal circuits and behavior.

In conclusion, our study illustrates the potential of partial reprogramming using YFs in both promoting neurogenesis and preventing neurodegeneration. When induced during development, increased neurogenesis leads to expansion of the neocortex and improved behavior. When induced in neurons at adult stages, YFs expression prevents the development of several AD-related hallmarks and cognitive decline. Hence, our findings unveil two novel applications for partial reprogramming and pave the way for further investigations related to neuronal rejuvenation.

## Author contributions

YS led the initial characterization of i4F-Rosa embryos, characterized the i4F-Nes mouse model and produced samples for mass spectrometry.

SZ led the mouse behavior work of i4F-Rosa and i4F:5xFAD models, performed IF experiments and Golgi stainings

XB contributed to the characterization of the i4F-Rosa embryos using different Dox treatments.

AS performed stereotaxic surgeries to transduce YFs in the 5xFAD mouse model.

CdC performed electron microscopy studies.

GSB quantified progenitor numbers and distribution in i4F-Nes embryos IB contributed to the Golgi stainings and analysis.

NA generated the first i4F-Rosa embryos treated with different Dox treatments JA oversaw the work using i4F-Rosa and i4F:5xFAD mouse models.

MA Provided the i4F mouse model and expertise to induce YFs *in vivo*.

MS oversaw the cell reprogramming aspects of the work and the induction of YFs *in vivo*.

RK oversaw the characterization of i4F-Rosa of i4F-Nes mouse models.

AG oversaw the mouse behaviour performed in adult i4F-Rosa, the work related to the 5xFAD mouse model and corresponding design of stereotaxic surgeries, IF and mouse behaviour.

DdT oversaw the work using i4F-Rosa and i4F-Nes mouse models, viral constructs, proteomics and cell biology aspects of the work.

All authors have contributed to the manuscript.

## Acknowledgements

We thank Ana López (María de Maeztu Unit of Excellence, Institute of Neurosciences, University of Barcelona, CEX2021-001158-M, Ministry of Science, Innovation and Universities) for technical support. We thank M. Calvo from the Advanced Microscopy service (CCiT, university of Barcelona) for help with confocal microscopy. We thank the proteomics facility of the Max Planck Institute of Biochemistry (Martinsried, Germany). We thank Patrick Hoffmann, Ekaterina Prilevskaya and Fady Shenouda for help with management of the animal colony. SZ was funded by the FPI fellowship from MINECO program. MA was funded by VHIO, Fero Foundation and Milky Way Research Foundation. MS was funded by the IRB, “laCaixa” Foundation, and by the Milky Way Research Foundation. This work was supported by grants from MCIN/AEI/10.13039/501100011033 and by European Union NextGenerationEU/PRTR: A.G.: PID2021-122258OB-I00 and CNS2022 – 135391 (“Consolidación” program). This work was also supported by the Max-Planck Society (to R.K.) and the Fundación Ramón Areces (CIVP21A7024) to A.G. DdT was funded by the Ramón y Cajal program (RYC-2017-23486) and MINECO project: RTI2018-095580-A-100 and PID2021-124852OB-100.

## Supplemental movie legends

**Supplemental movie S1: Cleared Whole-Mount i4F-Rosa and control embryonic brains at E15.5.** 3D reconstruction of embryonic brains treated with high-Dox (1mg/ml) for 4d (E10.5 to E14.5) and fixed at E15.5. Brains were cleared using a protocol that combined CUBIC and RIMS. After clearing, samples were stained with propidium iodide (5 mM) for 2 days to stain cell nuclei. Samples were mounted using RIMS buffer and imaged from dorsal to ventral at intervals of every 20µm. The movie displays sagittal sections of half a hemisphere for both the control (left) and i4F-Rosa (right). Additionally, the inset used for analyzing GZ thickness is depicted (Fig. 1B-C).

**Supplementary figure 1.**
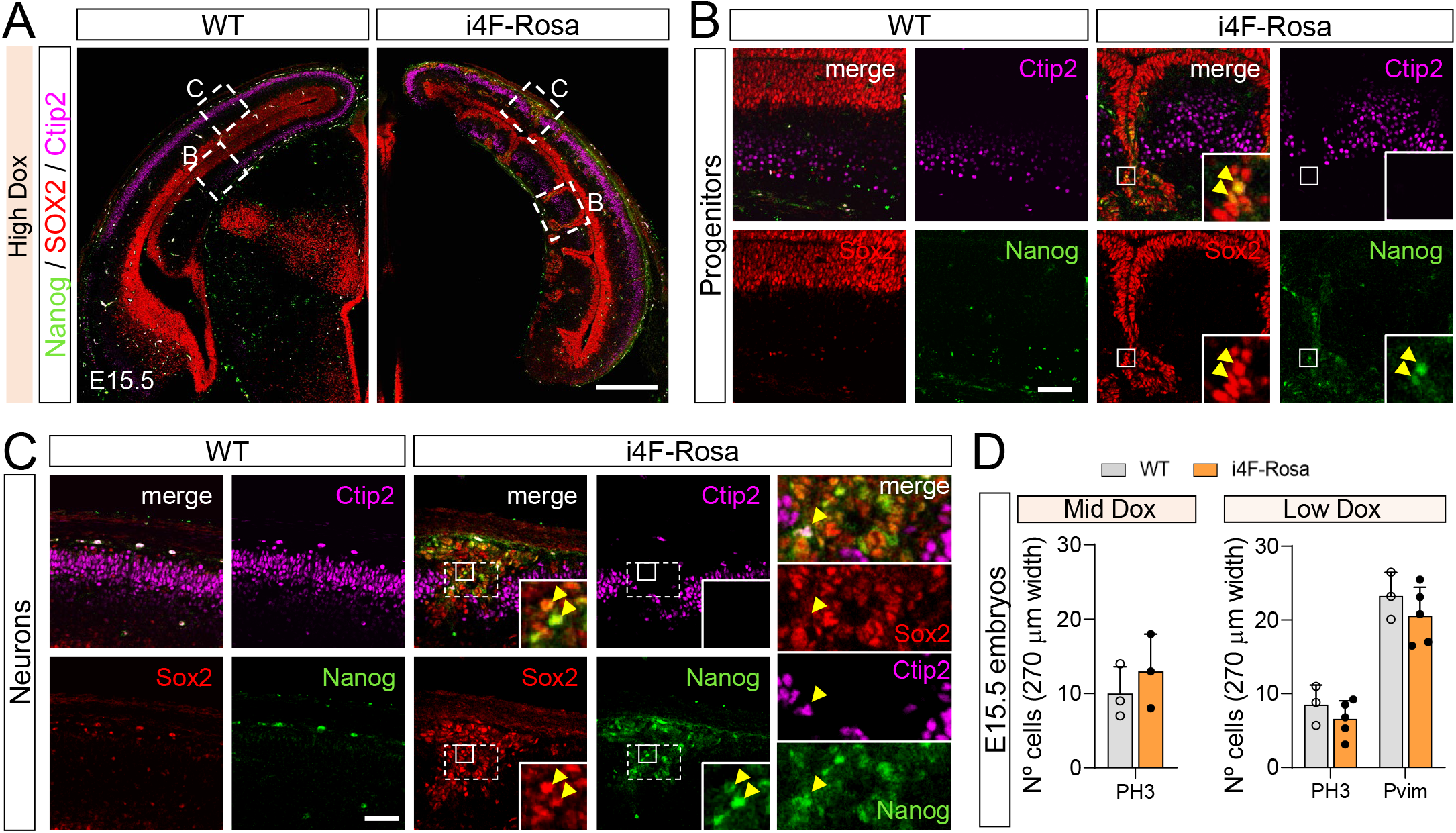
Strong YF induction alters progenitor and neuronal identity. (A) E15.5 coronal brain sections of WT and i4F-Rosa embryos, after 4d of high Dox, were stained for Nanog (an iPS cell marker, green), Sox2 (a neuronal progenitor marker and one of the YFs, red) and Ctip2 (neurons, magenta). Areas in dashed rectangles representing progenitors or neuron-enriched regionsare shown with higher magnification in (B) and (C), respectively. (B-C) Arrows in insets indicates double-positive Sox2/Nanog cells. Panel C has two insets indicating a Sox2/Nanog+ cell (small inset) and Ctip2/Nanog+ cell (inset on the right). (D) Quantification of data shown in (Fig. 1F and I). n=3-5 mice/group. The data are represented as mean ± SEM. Circles indicate values of individual brains.Scale bars, 500um (A) and 250um (B,C).

**Supplementary figure 2.**
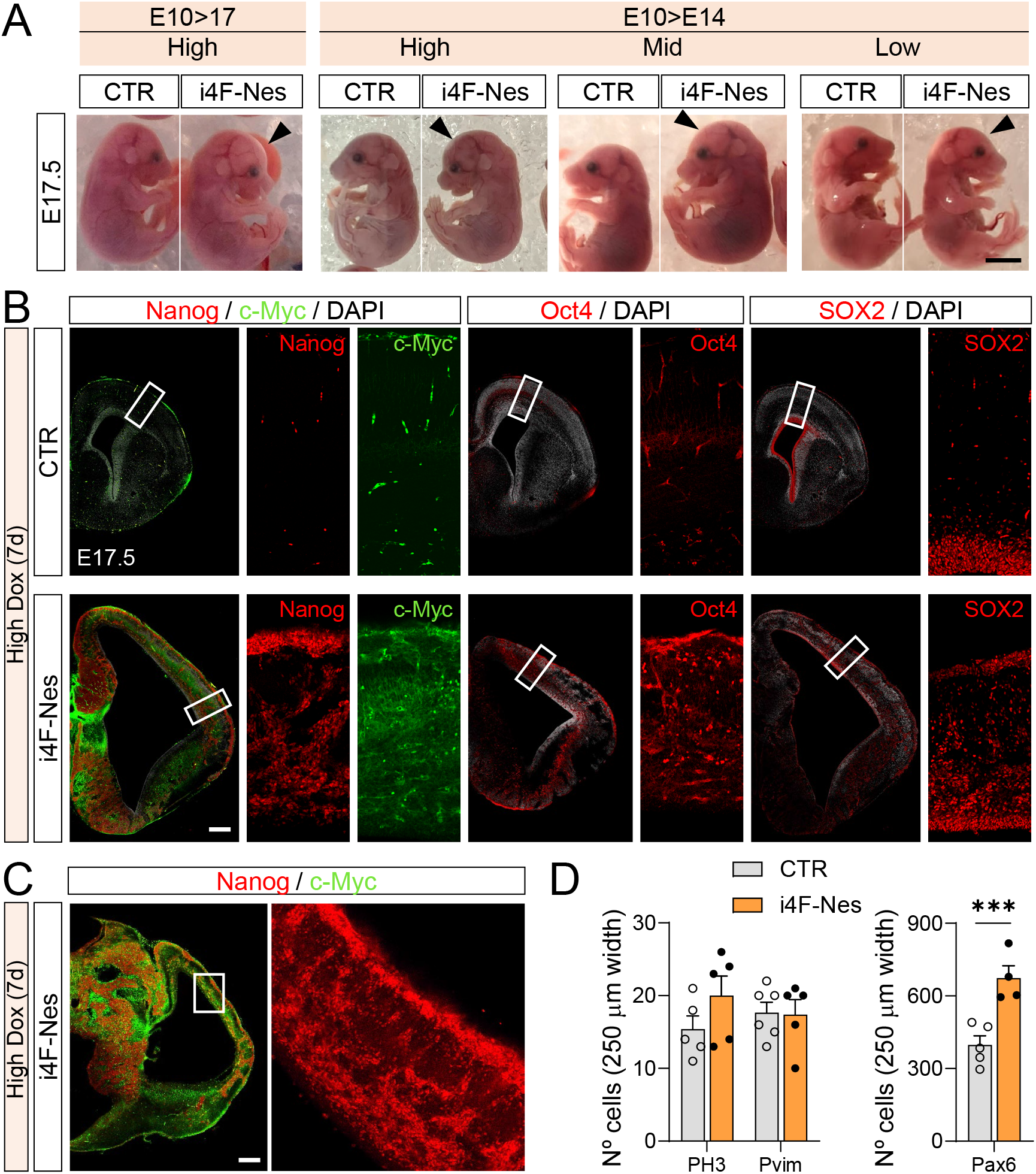
Effects of YF induction in brain size and cell identity. (A) CTR and i4F-Nes embryos at E17.5 with different Dox treatments. (B, C) E17.5 coronal brain sections, after 7d of high Dox, were stained for Nanog (an iPS cell marker, red), c-Myc (one of the YFs, red), Oct4 (one of the YFs, red) and Sox2 (a neuronal progenitor marker and one of the YFs, red) from left to right image. Areas in dashed rectangles are shown with higher magnification on the right. (D) Quantification of data shown in Figure 2D. n=4-6 mice/group. ***p<0.001, unpaired Student’s t test. Scale bars, 3mm (A) and 500um (B, C).

**Supplementary figure 3.**
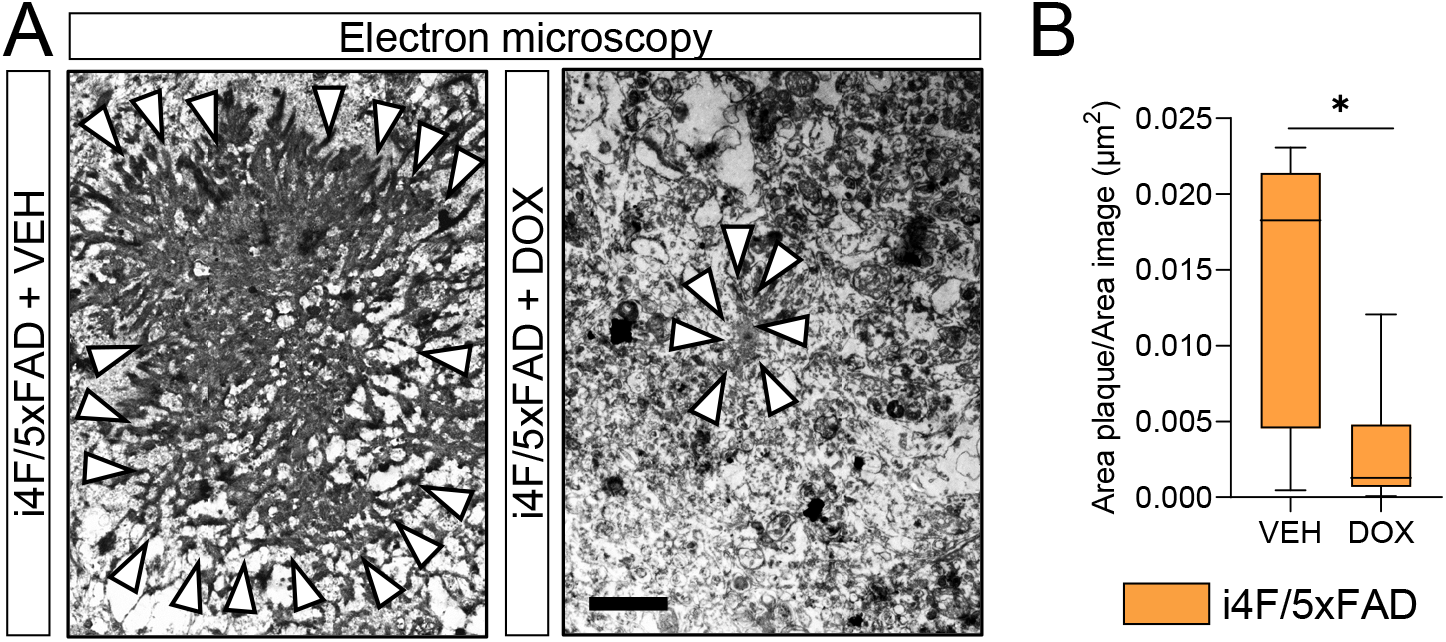
Effects of the YF induction in hippocampal plaques loading. (A) Amyloid plaque images obtained with electronic microscopy in the dentate gyrus (molecular layer). White arrows indicate the plaque boundary. (B) Quantification of Aβ plaques area in the dentate gyrus as in A. (t = 2.7434, df = 10; p = 0.0207). Unpaired Student’s t-test was employed in B (n = 6 per group). Data are mean ± SEM.

**Supplementary figure 4.**
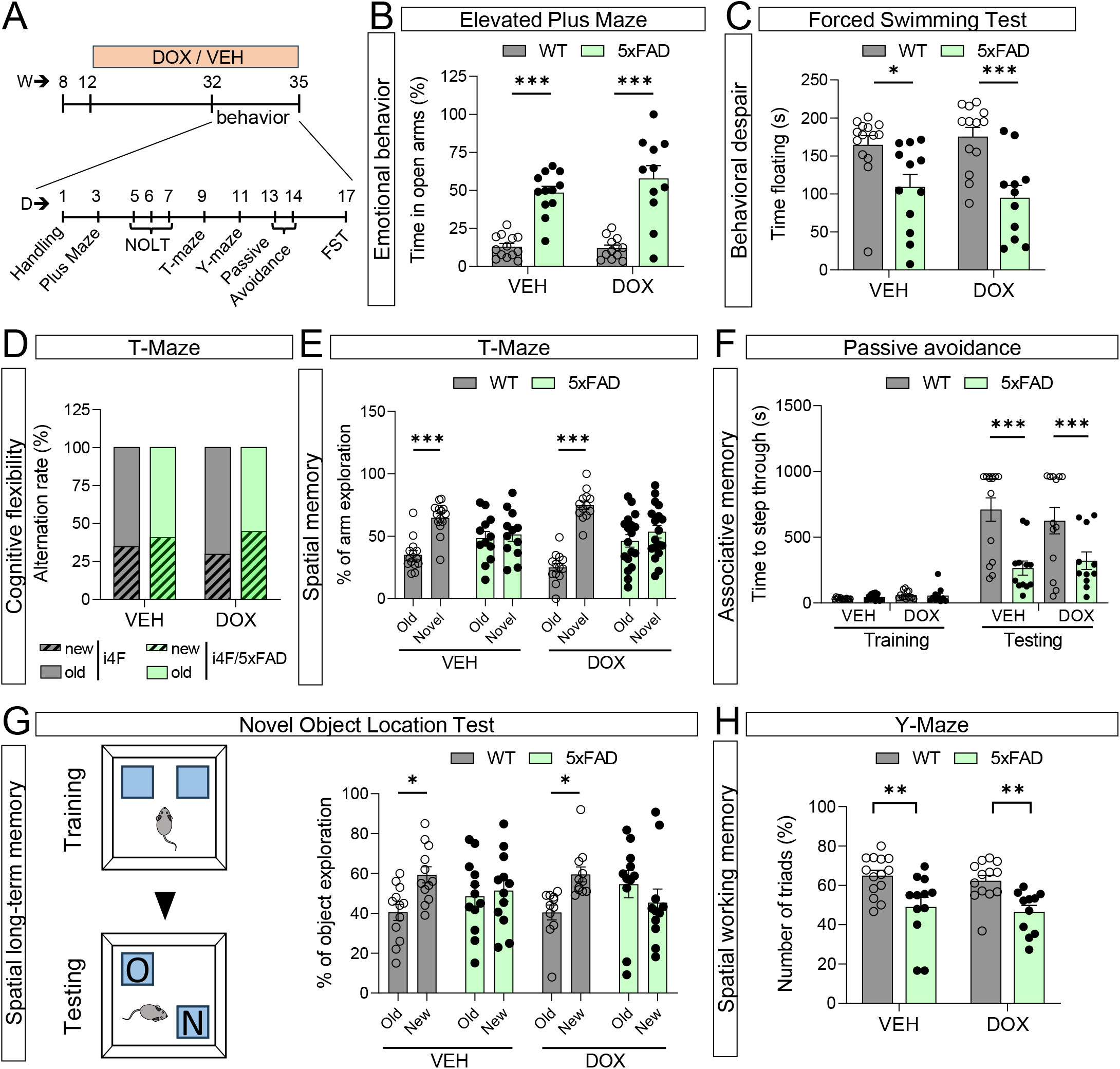
Doxycycline intermittent treatment does not impact mouse behavior. (A) Experimental design and sequence of behavioral tests performed. (B) Percentage of time spent in open arms in the elevated Plus Maze test. n=11-14 mice/group. ***p<0.001 respect to WT-VEH and WT-DOX groups respectively, Two-way ANOVA with Bonferroni’s post hoc test. (C) Time floating in the forced swimming test. n=11-14 mice/group. *p<0.05 and ***p<0.001 respect to WT-VEH and WT-DOX groups respectively, Two-way ANOVA with Bonferroni’s post hoc test. (D-E) Quantification of cognitive flexibility and Spatial memory in a T-maze. (D) Cognitive flexibility was measured as the first arm visited in the testing trial. Alternation is presented as %. n=11-14 mice/group. Pair comparisons were performed with the Chi-square test in. (E) Percentage of time exploring the novel arm (new context) versus the time exploring the old arm (old context). n=11-14 mice/group. ***P < 0.001 comparing % of novel arm vs old arm preference for each group. Two-way ANOVA with Bonferroni’s post hoc test. (F) Associative learning measured using the passive avoidance paradigm. Quantification of the latency (seconds) to step-through in both training and testing trial for all groups after receiving an electric shock (2 s/1 mA). n=11-14 mice/group. ***p<0.001 respect to WT-VEH and WT-DOX groups respectively. Two-way ANOVA with Bonferroni’s post hoc test. (G) Spatial long-term memory using the novel object location test. Percentage of time exploring the object placed in a new location (New) versus the time exploring the object placed in an old location (Old). n=11-14 mice/group. *p<0.05 comparing time exploring objects in new vs old location. Two-way ANOVA with Bonferroni’s post hoc test. (H) Quantification of spontaneous alternation (number of triads in %) in the Y-maze. n=11-14mice/group. **p<0.01 respect to 5xFAD-VEH and 5xFAD-DOX groups respectively. The data are represented as mean ± SEM. Circles indicate values of individual mice.

## STAR Methods

### Experimental model and subject details Animals

Experiments were performed during development (from E10.5 till E17.5) or adult mice (>8 weeks). The wild-type animals were from the C57BL/6N strain (Charles River labs, https://www.criver.com/). We used the i4F-A mouse line described previously to perform all the crosses^17^. For developmental studies, i4F-Rosa refers to i4F-A carrying rtTA within the ubiquitously-expressed Rosa26locus^17^. I4F-Nes was developed by crossing both i4F-A and rtTA with the nervous system-specific Nestin-Cre^84^. This floxed rtTA transgenic line [B6.Cg-Gt(ROSA)26Sor<tm1(rtTA,EGFP)Nagy>/J] was imported from Jackson Laboratories (Stock# 005670).

In studies related to neurodegeneration, we utilized the previously described 5xFAD mouse line^40^, which was bred with the i4F-A mouse line. The animals were housed with access to food and water ad libitum in a colony room kept at 19°C–22°C and 40%–60% humidity, under a 12:12 h light/dark cycle. In all studies, Doxycycline (BioChemica) was administered in the drinking water supplemented with 7.5% of sucrose as previously described^17^.

Experimental animals were used in accordance with the ethical guidelines (Declaration of Helsinki and NIH Publication no. 85-23, revised 1985, European Community Guidelines, and approved by the UB (CEEA: 55/21) and regional government (Generalitat de Catalunya: 11559) ethical committees. Animal experiments conducted at the MPI were performed following regulations from the government of Upper Bavaria. The animal study was reviewed and approved by the Regierung von Oberbayern under the license 55.2-2532.Vet_02-20-49.

### Experimental design using AAVs

We designed a viral construct to express rtTA only in principal neurons due to the AAV8 capsid and the Synapsin I promoter to drive the expression of the OSKM under the Tet-ON system. This pAAV8[TetOn]-TRE>ZsGreen1-rev(SYN1>tTS:T2A:rtTA virus was injected/transduced in the hippocampus of adult i-4F:WT and i-4F:5xFAD male mice at 8 weeks of age (see Fig 4A). At 12 weeks of age, all mice were treated with vehicle (VEH, sucrose 7.5%) or with doxycycline (Dox, 0.2 mg/kg in 7.5% sucrose) in drinking water 3 days/week. The treatment spanned 5 months. In the final month, when the mice were 8 months old, all were subjected to a comprehensive behavioral test battery (see Fig. 6 and behavioral assessments down below for further details). Last day treatment, all mice were sacrificed for subsequent histological and Golgi studies.

### Stereotaxic surgeries and viral constructs

Following anesthesia with ketamine/xylazine (100 and 10 mg/kg, respectively), we performed bilateral hippocampal injections of pAAV8[TetOn]-TRE>ZsGreen1-rev(SYN1>tTS:T2A:rtTA (2×1013 GC/ml in PBS, Vector Builder. Vector ID:VB210109-1009yyx). We used the following coordinates (millimeters) from bregma (anteroposterior and lateral) and from skull (dorsoventral); anteroposterior: −2.0; Lateral +/−1.25, and dorsoventral: −1.3 (CA1) and −2.1 (Dentate Gyrus). The cannula was left to deliver 1 μl of 1:1 virus in each depth during 2 min and five additional minutes were left to have complete virus diffusion. After 2 h of careful monitoring, mice were returned to their home cage for 4 weeks before starting the doxycycline (Dox) treatment.

### Behavioural assessments

#### Plus Maze

To analyze mouse anxiety, we used the elevated plus maze paradigm. Briefly, the plus maze was made of plastic and consisted of two opposing 30 × 8 cm open arms, and two opposing 30 × 8 cm arms enclosed by 15-cm-high walls. The maze was raised 50 cm from the floor and lit by dim light. Each mouse was placed in the central square of the raised plus maze, facing an open arm, and its behavior was scored for 5 min. At the end of each trial, any defecation was removed, and the apparatus was wiped with 30% ethanol. We recorded the time spent in the open arms, which normally correlates with low levels of anxiety. Animals were tracked and recorded with SMART junior software (Panlab).

#### Novel Object Location Test (NOLT)

The novel object location memory task evaluates spatial memory and is based on the ability of mice to recognize when a familiar object has been relocated. Exploration took place in an open-top arena with quadrangular form (45 × 45 cm). The light intensity was 40 lux throughout the arena. Mice were first habituated to the arena in the absence of objects (1 d, 30 min). Some distal cues were placed throughout the procedure. On the second day during the acquisition phase, mice could explore two duplicate objects (A1 and A2), which were placed close to the far corners of the arena for 10 min. After a delay of 24 h, one object was placed in the diagonally opposite corner. Thus, both objects in the phase were equally familiar, but one was in a new location. The position of the new object was counterbalanced between mice. Animals were tracked and recorded with SMART Junior software (Panlab).

#### Cognitive flexibility and memory in a T-maze

The apparatus was a wooden maze consisting of three arms, two of them situated at 180° from each other and the third situated at 90° with respect to the other ones representing the stem arm of the T. All three arms were 45 cm long, 8 cm wide and enclosed by a 20 cm wall. The maze was thoroughly painted with waterproof gray paint. Light intensity was 10-20 lux throughout the maze. Two identical guillotine doors provided entry in the arms situated at 180°. In the training trial, one arm was closed (novel arm) and mice were placed in the stem arm of the T (home arm) and allowed to explore this arm and the other available arm (familiar arm) for 10 min, after which they were returned to the home cage. After inter-trial interval of 1 h mice were placed in the stem arm of the T-maze and allowed to freely explore all three arms for 5 min (testing phase). The first choice to turn either to the familiar arm or to the new arm (alternation rate, %) and the time spent in the two arms situated at 180° (time in the new arm*100/total time in the two arms at 180°) were the two parameters evaluated in the testing phase.

#### Y-maze spontaneous alternation

The spontaneous alternation performance was tested using a transparent symmetrical Y-maze. Five objects of similar size (∼20 cm3) and highly perceptible were situated surrounding the maze at ∼15–20 cm outside the walls. Each mouse was placed in the center of the Y-maze and could explore freely through the maze during an 8 min session. The sequence and total number of arms entered were recorded. Arm entry was complete when the hind paws of the mouse had been completely placed in the arm. Percentage alternation is the number of triads containing entries into all three arms divided by the maximum possible alternations (dividing the number of alternations by number of possible triads x 100). As the reentry into the same arm was not counted for analysis, the chance performance level in this task was 50% in the choice between the arm mice visited more recently (non-alternation) and the other arm visited less recently (alternation).

#### Forced Swimming Test

The forced swimming test was used to evaluate behavioral despair. Animals were subjected to a 6 min trial during which they were forced to swim in an acrylic glass cylinder (35 cm of height × 20 cm of diameter) filled with water, and from which they could not escape. The time that the test animal spent in the cylinder without making any movements beyond those required to keep its head above water was measured.

#### Passive Avoidance Test

For the passive avoidance (light-dark) paradigm, we conducted the experiments in a 2-compartment box, where 1 compartment was dimly lit (20 lux) and preferable to a rodent and the other compartment was brightly lit (200 lux); both chambers were connected by a door (5 cm × 5 cm). During training, mice were placed into the aversive brightly lit compartment; and upon the entry into the preferred dimly lit compartment (with all 4 paws inside the dark chamber), mice were exposed to a mild foot shock (2 s foot shock, 1 mA intensity). The latency of mice to enter into the dark chamber was recorded. Twenty seconds after receiving the foot shock, mice were returned to the home cage until testing. After 24 h (long-term associative memory), animals were tested for retention. In the retention test, mice were returned to the brightly lit compartment again, and the latency to enter the shock paired compartment (dark chamber) was measured (retention or recall latency). Ten minutes was used as a time cutoff in the retention test. The animals that learned the task would avoid the location previously paired with the aversive stimulus and showed a greater latency to enter it.

#### Accelerating rotarod

Animals were placed on a motorized rod (30-mm diameter, Panlab). The rotation speed was gradually increased from 4 to 40 r.p.m. over the course of 5 min. The fall latency time was recorded when the animal was unable to keep up with the increasing speed and fell. Rotarod training/testing was performed four times per day with 30 min as intertrial time interval. The results show the average of fall latencies per trial during the 2 days of training.

#### Marble test

Briefly, mice were individually placed in a clear plastic box (35 x 20 x 15 cm) filled with approximately 5 cm depth of wood chip bedding lightly pressed to give a flat surface. Twenty 1.5 cm diameter glass marbles were placed on the surface, evenly spaced, each about 4 cm apart, so to form 5 rows of 4. The completely unburied (OUT), partially unburied (>50% OUT), partially buried (>50% IN) and completely buried (IN) marbles were manually quantified at the end of a 20-min test session.

#### Three chamber social interaction test

The apparatus (40 x 40 x 60 cm) consisted of three interconnected lined compartments with open doors. Subject mice were habituated to the apparatus for 5 min. After the habituation phase, the subjects were tested in the sociability task for 10 min. An unfamiliar mouse (stranger) was placed in one of the side chambers enclosed in a small, round wire cage that allowed nose contact between the bars but prevented fighting. A second, empty round wire cage, was placed in the opposite compartment. The subject mouse had a choice between the first, unfamiliar mouse (stranger) and the empty wire cage (empty). Time exploring each small cage were measured using the SMART junior software (Panlab).

### Golgi Staining and dendritic spine density analysis

Fresh brain right hemispheres were processed following the Golgi-Cox method as described previously^85^. Essentially, mouse brain hemispheres were incubated in the dark for 21 days in filtered dye solution (10 g L−1 K2Cr2O7, 10 g L−1 HgCl2, and 8 g L−1 K2CrO4). The tissue was then washed 3 × 2 min in water and 30 min in 90% ethanol (EtOH) (v/v); 200 μm sections were cut in 70% EtOH on a vibratome (Leica Microsystems) and washed in water for 5 min. Next, they were reduced in 16% (v/v) ammonia solution for 1 h before washing in water for 2 min and fixation in 10 g l−1 Na2S2O3 for 7 min. After a 2 min final wash in water, sections were mounted on superfrost coverslips, dehydrated for 3 min in 50%, then 70%, 80%, and 100% EtOH, incubated for 2 × 5 min in a 2:1 isopropanol:EtOH mixture, followed by 1 × 5 min in pure isopropanol and 2 × 5 min in xylol. Bright-field images of Golgi-impregnated stratum moleculare dendrites from hippocampal dentate gyrus granular neurons were captured with a Nikon DXM 1200F digital camera attached to a Nikon Eclipse E600 light microscope (100× oil objective). Only fully impregnated pyramidal neurons with their soma found entirely within the thickness of the section were used. Image z stacks were taken every 0.2 mm and at 1024 × 1024 pixel resolution, yielding an image with pixel dimensions of 49.25 × 49.25 mm. Z stacks were deconvolved using the Huygens software (Scientific Volume Imaging) to improve voxel resolution and to reduce optical aberration along the z axis. The total number of spines counting were performed by using the FIJI freeware (Wayne Rasband, NIH, RRID:SCR_003070)). At least 100-120 dendrites per group from at least 8-11 mice per group were counted. Picture acquisition and subsequent analysis were performed independently by two investigators blind to genotypes and results were then pooled.

### Immunohistochemistry

#### Adult samples

Mice were euthanized by cervical dislocation. Left hemispheres were removed and fixed for 72 h in 4% paraformaldehyde (PFA) in PBS. Thirty-micrometer coronal sections were obtained using a Leica vibratome (Leica VT1000S) and then cryoprotected until use. Serial coronal sections were then washed three times in PBS, permeabilized 15 min by shaking at room temperature with PBS containing (v/v) 0.3% Triton X-100 and 3% normal goat serum (Pierce Biotechnology). After three washes, brain sections were incubated overnight by shaking at 4°C with antibodies for anti-Aβ (Mouse 1:500, 218111, clone NT78, Synaptic Systems. RRID:AB_11041707), anti-NeuN (Mouse 1:1000, MAB377, Chemicon. RRID:AB_2298772) and anti-Pyk2 (Rabbit, 1:500, P3902. Sigma. RRID:AB_261041) in PBS with 0.2 g/L sodium azide. After incubation with primary antibody, sections were washed three times and then placed 2 h on a shaking incubator at room temperature with the subtype-specific fluorescent secondary AlexaFluor-488 anti-rabbit (1:250, Thermo Fisher Scientific catalog #A32731, RRID:AB_2633280) or anti-mouse 555 (1:250, Thermo Fisher Scientific catalog #A32727, RRID:AB_2633276). No signal was detected in control sections incubated in the absence of the primary antibody. Images were acquired using a Zeiss LSM880 confocal laser scanning microscope or SP8 laser scanning confocal spectral microscope (Leica Microsystems, Germany). Images were taken using a 20× numerical aperture objective and 2 Airy disk pinhole and processed with ImageJ^86^ (v1.53f) or Cell Profiler^87^ (v.2) software.

#### Embryonic samples

Embryonic brains were fixed in 4% PFA over-night. Vibratome sections (50-70um thick) were incubated with primary antibodies after 2 hour of permeabilization and blocking with 1% BSA, 0.3% Triton X-100/PBS, 5% donkey serum (Pierce Biotechnology). We used rabbit anti-Satb2 antibody 1/300 (Abcam), rat anti-Ctip2 1/300 (Abcam), rabbit anti-Pax6 1/300 (BioLegend), goat anti-Sox2 1/300 (R&D), rabbit anti-Histone H3 1/300 (Abcam), mouse anti-Pvim 1/300 (Abcam). The secondary antibodies were Alexa Fluor 488-, 555- and 647-conjugated goat or donkey anti-rabbit/mouse/goat (Molecular Probes 1:400). Images were acquired using a Zeiss LSM880 confocal laser scanning microscope or SP8 laser scanning confocal spectral microscope (Leica Microsystems, Germany). Images were taken using a 20× numerical aperture objective and 2 Airy disk pinhole and processed with ImageJ^86^ (v1.53f) or Cell Profiler^87^ (v.2) software.

### Electron microscope experiments

As we previously described^81^, mice were transcardially perfused with a solution containing 4% PFA and 0.1% glutaraldehyde made up of 0.1 m PB, pH 7.4. Brains were then immersed in the same fixative for 12 h at 4°C. Tissue blocks containing the hippocampus were dissected and washed in 0.1 m PB, cryoprotected in 100 and 200 g/L sucrose in 0.1 m PB, and freeze-thawed in isopentane and liquid nitrogen. Samples were postfixed in 2.5% glutaraldehyde made up of 0.1 m phosphate buffer for 20 min, washed and treated with 2% osmium tetroxide in PB for 20 min. They were dehydrated in a series of ethanol and flat embedded in epoxy resin (EPON 812 Polysciences). After polymerization, blocks from the dentate gyrus (DG) region were cut at 70 nm thickness using an ultramicrotome (Ultracut E Leica Microsystems). Sections were cut with a diamond knife, picked up on formvar-coated 200 mesh nickel grids. For etching resin and remove osmium, sections were treated with saturated aqueous sodium periodate (NaIO4). They were then observed with a CM-100 electron microscope (Philips). Digital images were obtained with a CCD camera (Gatan Orius). In ultrathin sections, the density of synaptic vesicles in the molecular layer was calculated by counting the number of vesicles within a defined presynaptic area. The area of postsynaptic densities located in the molecular layer was also evaluated. All these calculations were performed using the ImageJ.

### Mass spectrometry experiments

Fresh E15.5 mouse cortices or dentate gyrus from adult mice were homogenized for 1min at 4°C with an electric homogenizer using the following lysis buffer: 50 mM Tris-HCL (pH 7.4), 150mM NaCl, 2mM EDTA, 1% Triton X-100 and protease inhibitors (Roche ref. 04693116001). Samples were incubated on ice for 20 min and centrifuged for 10 min at 3000 rpm. Supernatant was collected and protein was measured using the Bio-Rad protein assay (Biorad, 5000001). 25-50 μl wof protein at a final concentration of 1-2 μg/μl in lysis buffer were processed for mass spectrometry (MaxQuant run, Proteomic facility, Max Planck Institute of Biochemistry, Martinsried, Germany). 4-6 independent samples per processed per condition in all experiments. Analysis of differentially expressed proteins and volcano plots were generated using the DEP package in R-studio.

### Cleared whole-mount embryonic brain

Dissected embryonic mouse brains were cleared by combining the protocols CUBIC^88^ and RIMS^89^. Briefly, brains were fixed in 4% PFA overnight. After washing with PBS, they were immersed in CUBIC solution (25% Urea and 25% tetrakis in water) with gentle shaking till transparency at 37°. Brains were washed in PBS for 2d and post-fixed in 4% PFA for 2h. They were then stained with 5mM propidium iodine dissolved in PBS for 2d. Following another 1d PBS wash, samples were mounted in RIMS (76% Histodenz-0.02M PB-0.01%SodiumAzide-0.1%Tween-20) for confocal acquisition. Images were acquired using a Zeiss LSM880 confocal laser scanning microscope using a 10× numerical aperture objective and 2.5 Airy disk pinhole and processed with Imaris and ImageJ software.

### Quantification and Statistical Analysis

Statistical analyses were performed using GraphPad Prism v.8 (GraphPad Software, La Jolla, California, USA). A two-tailed unpaired Student’s t-test was employed when comparing two groups, while one-way ANOVA with Tukey’s post hoc analysis or two-way ANOVA with Bonferroni’s post hoc test was used for multiple group comparisons, as appropriate. Normal distribution was tested with d’Agostino and Pearson omnibus test. In case of no normal distribution corrections were applied (Mann-Whitney or Dunn’s test). P values represent ∗p ≤ 0.05, ∗∗p ≤ 0.01, ∗∗∗p ≤ 0.001 and ∗∗∗∗p ≤ 0.0001. All data are presented as the mean ± s.e.m, whisker plots or dot plots. All sample sizes and definitions are provided in the figure legends. All experiments in this study were conducted in a blinded and randomized manner. All mice bred for the experiments were used for preplanned experiments and randomized to experimental groups.

## References

1. Takahashi, K., and Yamanaka, S. (2006). Induction of pluripotent stem cells from mouse embryonic and adult fibroblast cultures by defined factors. Cell 126, 663–676. 10.1016/J.CELL.2006.07.024.

2. Ocampo, A., Reddy, P., Martinez-Redondo, P., Platero-Luengo, A., Hatanaka, F., Hishida, T., Li, M., Lam, D., Kurita, M., Beyret, E., et al. (2016). In Vivo Amelioration of Age-Associated Hallmarks by Partial Reprogramming. Cell 167, 1719–1733.e12. 10.1016/j.cell.2016.11.052.

3. Chondronasiou, D., Gill, D., Mosteiro, L., Urdinguio, R.G., Berenguer-Llergo, A., Aguilera, M., Durand, S., Aprahamian, F., Nirmalathasan, N., Abad, M., et al. (2022). Multi-omic rejuvenation of naturally aged tissues by a single cycle of transient reprogramming. Aging Cell 21. 10.1111/ACEL.13578.

4. Sarkar, T.J., Quarta, M., Mukherjee, S., Colville, A., Paine, P., Doan, L., Tran, C.M., Chu, C.R., Horvath, S., Qi, L.S., et al. (2020). Transient non-integrative expression of nuclear reprogramming factors promotes multifaceted amelioration of aging in human cells. Nat Commun 11. 10.1038/S41467-020-15174-3.

5. Lu, Y., Brommer, B., Tian, X., Krishnan, A., Meer, M., Wang, C., Vera, D.L., Zeng, Q., Yu, D., Bonkowski, M.S., et al. (2020). Reprogramming to recover youthful epigenetic information and restore vision. Nature 588, 124–129. 10.1038/S41586-020-2975-4.

6. Yang, J.H., Hayano, M., Griffin, P.T., Amorim, J.A., Bonkowski, M.S., Apostolides, J.K., Salfati, E.L., Blanchette, M., Munding, E.M., Bhakta, M., et al. (2023). Loss of epigenetic information as a cause of mammalian aging. Cell 186, 305–326.e27. 10.1016/J.CELL.2022.12.027.

7. Zhang, W., Li, J., Suzuki, K., Qu, J., Wang, P., Zhou, J., Liu, X., Ren, R., Xu, X., Ocampo, A., et al. (2015). Aging stem cells. A Werner syndrome stem cell model unveils heterochromatin alterations as a driver of human aging. Science 348, 1160–1163. 10.1126/SCIENCE.AAA1356.

8. Mahmoudi, S., and Brunet, A. (2012). Aging and reprogramming: a two-way street. Curr Opin Cell Biol 24, 744–756. 10.1016/J.CEB.2012.10.004.

9. Browder, K.C., Reddy, P., Yamamoto, M., Haghani, A., Guillen, I.G., Sahu, S., Wang, C., Luque, Y., Prieto, J., Shi, L., et al. (2022). In vivo partial reprogramming alters age-associated molecular changes during physiological aging in mice. Nature Aging 2022 2:3 2, 243–253. 10.1038/s43587-022-00183-2.

10. Hishida, T., Yamamoto, M., Hishida-Nozaki, Y., Shao, C., Huang, L., Wang, C., Shojima, K., Xue, Y., Hang, Y., Shokhirev, M., et al. (2022). In vivo partial cellular reprogramming enhances liver plasticity and regeneration. Cell Rep 39, 110730. 10.1016/J.CELREP.2022.110730.

11. Wang, C., Rabadan Ros, R., Martinez-Redondo, P., Ma, Z., Shi, L., Xue, Y., Guillen-Guillen, I., Huang, L., Hishida, T., Liao, H.K., et al. (2021). In vivo partial reprogramming of myofibers promotes muscle regeneration by remodeling the stem cell niche. Nat Commun 12. 10.1038/S41467-021-23353-Z.

12. Farber, G., Liu, J., and Qian, L. (2022). OSKM-mediated reversible reprogramming of cardiomyocytes regenerates injured myocardium. Cell Regen 11. 10.1186/S13619-021-00106-3.

13. Chen, Y., Lüttmann, F.F., Schoger, E., Schöler, H.R., Zelarayán, L.C., Kim, K.P., Haigh, J.J., Kim, J., and Braun, T. (2021). Reversible reprogramming of cardiomyocytes to a fetal state drives heart regeneration in mice. Science 373, 1537–1540. 10.1126/SCIENCE.ABG5159.

14. Chondronasiou, D., Martinez de Villarreal, J., Melendez, E., Lynch, C.J., Pozo, N. del, Kovatcheva, M., Aguilera, M., Prats, N., Real, F.X., and Serrano, M. (2022). Deciphering the roadmap of in vivo reprogramming toward pluripotency. Stem Cell Reports 17, 2501–2517. 10.1016/J.STEMCR.2022.09.009.

15. Tran, K.A., Pietrzak, S.J., Zaidan, N.Z., Siahpirani, A.F., McCalla, S.G., Zhou, A.S., Iyer, G., Roy, S., and Sridharan, R. (2019). Defining Reprogramming Checkpoints from Single-Cell Analyses of Induced Pluripotency. Cell Rep 27, 1726–1741.e5. 10.1016/J.CELREP.2019.04.056.

16. Maza, I., Caspi, I., Zviran, A., Chomsky, E., Rais, Y., Viukov, S., Geula, S., Buenrostro, J.D., Weinberger, L., Krupalnik, V., et al. (2015). Transient acquisition of pluripotency during somatic cell transdifferentiation with iPSC reprogramming factors. Nat Biotechnol 33, 769–774. 10.1038/NBT.3270.

17. Abad, M., Mosteiro, L., Pantoja, C., Cañamero, M., Rayon, T., Ors, I., Graña, O., Megías, D., Domínguez, O., Martinez, D., et al. (2013). Reprogramming in vivo produces teratomas and iPS cells with totipotency features. Nature 502, 340–345. 10.1038/nature12586.

18. Xia, X., Jiang, Q., McDermott, J., and Han, J.D.J. (2018). Aging and Alzheimer’s disease: Comparison and associations from molecular to system level. Aging Cell 17. 10.1111/ACEL.12802.

19. Cai, H., Cong, W., Ji, S., Rothman, S., Maudsley, S., and Martin, B. (2012). Metabolic Dysfunction in Alzheimers Disease and Related Neurodegenerative Disorders. Curr Alzheimer Res 9, 5–17. 10.2174/156720512799015064.

20. HUANG, W.-J., ZHANG, X., and CHEN, W.-W. (2016). Role of oxidative stress in Alzheimer’s disease. Biomed Rep 4, 519–522. 10.3892/br.2016.630.

21. Reza-Zaldivar, E.E., Hernández-Sápiens, M.A., Minjarez, B., Gómez-Pinedo, U., Sánchez-González, V.J., Márquez-Aguirre, A.L., and Canales-Aguirre, A.A. (2020). Dendritic Spine and Synaptic Plasticity in Alzheimer’s Disease: A Focus on MicroRNA. Front Cell Dev Biol 8. 10.3389/fcell.2020.00255.

22. Yang, S.G., Wang, X.W., Qian, C., and Zhou, F.Q. (2022). Reprogramming neurons for regeneration: The fountain of youth. Prog Neurobiol 214, 102284. 10.1016/J.PNEUROBIO.2022.102284.

23. Valadez-Barba, V., Cota-Coronado, A., Hernández-Pérez, O.R., Lugo-Fabres, P.H., Padilla-Camberos, E., Díaz, N.F., and Díaz-Martinez, N.E. (2020). iPSC for modeling neurodegenerative disorders. Regen Ther 15, 332. 10.1016/J.RETH.2020.11.006.

24. Brot, S., Thamrin, N.P., Bonnet, M.L., Francheteau, M., Patrigeon, M., Belnoue, L., and Gaillard, A. (2022). Long-Term Evaluation of Intranigral Transplantation of Human iPSC-Derived Dopamine Neurons in a Parkinson’s Disease Mouse Model. Cells 11. 10.3390/CELLS11101596.

25. Han, F., Wang, W., Chen, B., Chen, C., Li, S., Lu, X., Duan, J., Zhang, Y., Zhang, Y.A., Guo, W., et al. (2015). Human induced pluripotent stem cell-derived neurons improve motor asymmetry in a 6-hydroxydopamine-induced rat model of Parkinson’s disease. Cytotherapy 17, 665–679. 10.1016/J.JCYT.2015.02.001.

26. Rodríguez-Matellán, A., Alcazar, N., Hernández, F., Serrano, M., and Ávila, J. (2020). In Vivo Reprogramming Ameliorates Aging Features in Dentate Gyrus Cells and Improves Memory in Mice. Stem Cell Reports 15, 1056–1066. 10.1016/j.stemcr.2020.09.010.

27. Zalc, A., Sinha, R., Gulati, G.S., Wesche, D.J., Daszczuk, P., Swigut, T., Weissman, I.L., and Wysocka, J. (2021). Reactivation of the pluripotency program precedes formation of the cranial neural crest. Science (1979) 371. 10.1126/science.abb4776.

28. Ohnishi, K., Semi, K., Yamamoto, T., Shimizu, M., Tanaka, A., Mitsunaga, K., Okita, K., Osafune, K., Arioka, Y., Maeda, T., et al. (2014). Premature termination of reprogramming in vivo leads to cancer development through altered epigenetic regulation. Cell 156, 663–677. 10.1016/j.cell.2014.01.005.

29. Yeo, R.W., Zhou, O.Y., Zhong, B.L., Sun, E.D., Navarro Negredo, P., Nair, S., Sharmin, M., Ruetz, T.J., Wilson, M., Kundaje, A., et al. (2023). Chromatin accessibility dynamics of neurogenic niche cells reveal defects in neural stem cell adhesion and migration during aging. Nature Aging 2023 3:7 3, 866–893. 10.1038/s43587-023-00449-3.

30. Galakhova, A.A., Hunt, S., Wilbers, R., Heyer, D.B., de Kock, C.P.J., Mansvelder, H.D., and Goriounova, N.A. (2022). Evolution of cortical neurons supporting human cognition. Trends Cogn Sci 26, 909–922. 10.1016/J.TICS.2022.08.012.

31. Fernández, V., Llinares-Benadero, C., and Borrell, V. (2016). Cerebral cortex expansion and folding: what have we learned? EMBO J 35, 1021–1044. 10.15252/embj.201593701.

32. Lee, C., Kim, Y., and Kaang, B.K. (2022). The Primary Motor Cortex: The Hub of Motor Learning in Rodents. Neuroscience 485, 163–170. 10.1016/J.NEUROSCIENCE.2022.01.009.

33. Levy, D.R., Tamir, T., Kaufman, M., Parabucki, A., Weissbrod, A., Schneidman, E., and Yizhar, O. (2019). Dynamics of social representation in the mouse prefrontal cortex. Nature Neuroscience 2019 22:12 22, 2013–2022. 10.1038/s41593-019-0531-z.

34. Shiotsuki, H., Yoshimi, K., Shimo, Y., Funayama, M., Takamatsu, Y., Ikeda, K., Takahashi, R., Kitazawa, S., and Hattori, N. (2010). A rotarod test for evaluation of motor skill learning. J Neurosci Methods 189, 180–185. 10.1016/J.JNEUMETH.2010.03.026.

35. Kedia, S., and Chattarji, S. (2014). Marble burying as a test of the delayed anxiogenic effects of acute immobilisation stress in mice. J Neurosci Methods 233, 150–154. 10.1016/J.JNEUMETH.2014.06.012.

36. Yang, M., Silverman, J.L., and Crawley, J.N. (2011). Automated Three-Chambered Social Approach Task for Mice. Curr Protoc Neurosci 56, 8.26.1-8.26.16. 10.1002/0471142301.NS0826S56.

37. Browder, K.C., Reddy, P., Yamamoto, M., Haghani, A., Guillen, I.G., Sahu, S., Wang, C., Luque, Y., Prieto, J., Shi, L., et al. (2022). In vivo partial reprogramming alters age-associated molecular changes during physiological aging in mice. Nature Aging 2022 2:3 2, 243–253. 10.1038/s43587-022-00183-2.

38. Hishida, T., Yamamoto, M., Hishida-Nozaki, Y., Shao, C., Huang, L., Wang, C., Shojima, K., Xue, Y., Hang, Y., Shokhirev, M., et al. (2022). In vivo partial cellular reprogramming enhances liver plasticity and regeneration. Cell Rep 39, 110730. 10.1016/J.CELREP.2022.110730.

39. Lu, Y., Brommer, B., Tian, X., Krishnan, A., Meer, M., Wang, C., Vera, D.L., Zeng, Q., Yu, D., Bonkowski, M.S., et al. (2020). Reprogramming to recover youthful epigenetic information and restore vision. Nature 588, 124–129. 10.1038/S41586-020-2975-4.

40. Oakley, H., Cole, S.L., Logan, S., Maus, E., Shao, P., Craft, J., Guillozet-Bongaarts, A., Ohno, M., Disterhoft, J., Van Eldik, L., et al. (2006). Intraneuronal beta-amyloid aggregates, neurodegeneration, and neuron loss in transgenic mice with five familial Alzheimer’s disease mutations: potential factors in amyloid plaque formation. J Neurosci 26, 10129–10140. 10.1523/JNEUROSCI.1202-06.2006.

41. De Pins, B., Cifuentes-Díaz, C., Thamila Farah, A., López-Molina, L., Montalban, E., Sancho-Balsells, A., López, A., Ginés, S., Delgado-García, J.M., Alberch, J., et al. (2019). Conditional BDNF Delivery from Astrocytes Rescues Memory Deficits, Spine Density, and Synaptic Properties in the 5xFAD Mouse Model of Alzheimer Disease. J Neurosci 39, 2441–2458. 10.1523/JNEUROSCI.2121-18.2019.

42. Kim, D.K., Han, D., Park, J., Choi, H., Park, J.C., Cha, M.Y., Woo, J., Byun, M.S., Lee, D.Y., Kim, Y., et al. (2019). Deep proteome profiling of the hippocampus in the 5XFAD mouse model reveals biological process alterations and a novel biomarker of Alzheimer’s disease. Exp Mol Med 51. 10.1038/S12276-019-0326-Z.

43. Babcock, K.R., Page, J.S., Fallon, J.R., and Webb, A.E. (2021). Adult Hippocampal Neurogenesis in Aging and Alzheimer’s Disease. Stem Cell Reports 16, 681–693. 10.1016/J.STEMCR.2021.01.019.

44. Ge, S.X., Jung, D., Jung, D., and Yao, R. (2020). ShinyGO: a graphical gene-set enrichment tool for animals and plants. Bioinformatics 36, 2628–2629. 10.1093/BIOINFORMATICS/BTZ931.

45. Shannon, P., Markiel, A., Ozier, O., Baliga, N.S., Wang, J.T., Ramage, D., Amin, N., Schwikowski, B., and Ideker, T. (2003). Cytoscape: a software environment for integrated models of biomolecular interaction networks. Genome Res 13, 2498–2504. 10.1101/GR.1239303.

46. Griñán-Ferré, C., Sarroca, S., Ivanova, A., Puigoriol-Illamola, D., Aguado, F., Camins, A., Sanfeliu, C., and Pallàs, M. (2016). Epigenetic mechanisms underlying cognitive impairment and Alzheimer disease hallmarks in 5XFAD mice. Aging 8, 664–684. 10.18632/AGING.100906.

47. Schneider, F., Baldauf, K., Wetzel, W., and Reymann, K.G. (2015). Effects of methylphenidate on the behavior of male 5xFAD mice. Pharmacol Biochem Behav 128, 68–77. 10.1016/J.PBB.2014.11.006.

48. Devi, L., and Ohno, M. (2015). TrkB reduction exacerbates Alzheimer’s disease-like signaling aberrations and memory deficits without affecting β-amyloidosis in 5XFAD mice. Transl Psychiatry 5. 10.1038/TP.2015.55.

49. Fanselow, M.S., and Dong, H.W. (2010). Are The Dorsal and Ventral Hippocampus functionally distinct structures? Neuron 65, 7. 10.1016/J.NEURON.2009.11.031.

50. Lalonde, R. (2002). The neurobiological basis of spontaneous alternation. Neurosci Biobehav Rev 26, 91–104. 10.1016/S0149-7634(01)00041-0.

51. Singh, P.B., and Zhakupova, A. (2022). Age reprogramming: cell rejuvenation by partial reprogramming. Development (Cambridge) 149. 10.1242/dev.200755.

52. Manukyan, M., and Singh, P.B. (2012). Epigenetic rejuvenation. Genes Cells 17, 337–343. 10.1111/J.1365-2443.2012.01595.X.

53. Gill, D., Parry, A., Santos, F., Okkenhaug, H., Todd, C.D., Hernando-Herraez, I., Stubbs, T.M., Milagre, I., and Reik, W. (2022). Multi-omic rejuvenation of human cells by maturation phase transient reprogramming. Elife 11. 10.7554/ELIFE.71624.

54. Roux, A.E., Zhang, C., Paw, J., Zavala-Solorio, J., Malahias, E., Vijay, T., Kolumam, G., Kenyon, C., and Kimmel, J.C. (2022). Diverse partial reprogramming strategies restore youthful gene expression and transiently suppress cell identity. Cell Syst 13, 574–587.e11. 10.1016/j.cels.2022.05.002.

55. Liu, X., Huang, J., Chen, T., Wang, Y., Xin, S., Li, J., Pei, G., and Kang, J. (2008). Yamanaka factors critically regulate the developmental signaling network in mouse embryonic stem cells. Cell Research 2008 18:12 18, 1177–1189. 10.1038/cr.2008.309.

56. Sridharan, R., Tchieu, J., Mason, M.J., Yachechko, R., Kuoy, E., Horvath, S., Zhou, Q., and Plath, K. (2009). Role of the murine reprogramming factors in the induction of pluripotency. Cell 136, 364– 377. 10.1016/J.CELL.2009.01.001.

57. Memi, F., Killen, A.C., Barber, M., Parnavelas, J.G., and Andrews, W.D. (2019). Cadherin 8 regulates proliferation of cortical interneuron progenitors. Brain Struct Funct 224, 277–292. 10.1007/S00429-018-1772-4/FIGURES/7.

58. Kam, J.W.K., Dumontier, E., Baim, C., Brignall, A.C., da Silva, D.M., Cowan, M., Kennedy, T.E., and Cloutier, J.F. (2016). RGMB and neogenin control cell differentiation in the developing olfactory epithelium. Development 143, 1534–1546. 10.1242/DEV.118638.

59. Llinares-Benadero, C., and Borrell, V. (2019). Deconstructing cortical folding: genetic, cellular and mechanical determinants. Nat Rev Neurosci 20, 161–176. 10.1038/s41583-018-0112-2.

60. Zhao, J., Feng, C., Wang, W., Su, L., and Jiao, J. (2022). Human SERPINA3 induces neocortical folding and improves cognitive ability in mice. Cell Discov 8. 10.1038/S41421-022-00469-0.

61. Xing, L., Kubik-Zahorodna, A., Namba, T., Pinson, A., Florio, M., Prochazka, J., Sarov, M., Sedlacek, R., and Huttner, W.B. (2021). Expression of human-specific ARHGAP11B in mice leads to neocortex expansion and increased memory flexibility. EMBO J 40. 10.15252/EMBJ.2020107093.

62. Buitrago, M.M., Schulz, J.B., Dichgans, J., and Luft, A.R. (2004). Short and long-term motor skill learning in an accelerated rotarod training paradigm. Neurobiol Learn Mem 81, 211–216. 10.1016/J.NLM.2004.01.001.

63. Kawai, R., Markman, T., Poddar, R., Ko, R., Fantana, A.L., Dhawale, A.K., Kampff, A.R., and Ölveczky, B.P. (2015). Motor cortex is required for learning but not executing a motor skill. Neuron 86, 800. 10.1016/J.NEURON.2015.03.024.

64. Bicks, L.K., Koike, H., Akbarian, S., and Morishita, H. (2015). Prefrontal cortex and social cognition in mouse and man. Front Psychol 6, 166005. 10.3389/FPSYG.2015.01805/BIBTEX.

65. Smaers, J.B., Gómez-Robles, A., Parks, A.N., and Sherwood, C.C. (2017). Exceptional Evolutionary Expansion of Prefrontal Cortex in Great Apes and Humans. Current Biology 27, 714–720. 10.1016/J.CUB.2017.01.020.

66. Kamil, A.C. (2004). Sociality and the evolution of intelligence. Trends Cogn Sci 8, 195–197. 10.1016/j.tics.2004.03.002.

67. Doeser, M.C., Schöler, H.R., and Wu, G. (2018). Reduction of Fibrosis and Scar Formation by Partial Reprogramming In Vivo. Stem Cells 36, 1216–1225. 10.1002/STEM.2842.

68. Dorostkar, M.M., Zou, C., Blazquez-Llorca, L., and Herms, J. (2015). Analyzing dendritic spine pathology in Alzheimer’s disease: problems and opportunities. Acta Neuropathol 130. 10.1007/S00401-015-1449-5.

69. Yuki, D., Sugiura, Y., Zaima, N., Akatsu, H., Takei, S., Yao, I., Maesako, M., Kinoshita, A., Yamamoto, T., Kon, R., et al. (2014). DHA-PC and PSD-95 decrease after loss of synaptophysin and before neuronal loss in patients with Alzheimer’s disease. Sci Rep 4. 10.1038/SREP07130.

70. Crowe, S.E., and Ellis-Davies, G.C.R. (2014). Spine pruning in 5xFAD mice starts on basal dendrites of layer 5 pyramidal neurons. Brain Struct Funct 219, 571–580. 10.1007/S00429-013-0518-6.

71. Hongpaisan, J., Sun, M.K., and Alkon, D.L. (2011). PKC ε activation prevents synaptic loss, Aβ elevation, and cognitive deficits in Alzheimer’s disease transgenic mice. J Neurosci 31, 630–643. 10.1523/JNEUROSCI.5209-10.2011.

72. Brigman, J.L., Wright, T., Talani, G., Prasad-Mulcare, S., Jinde, S., Seabold, G.K., Mathur, P., Davis, M.I., Bock, R., Gustin, R.M., et al. (2010). Loss of GluN2B-containing NMDA receptors in CA1 hippocampus and cortex impairs long-term depression, reduces dendritic spine density, and disrupts learning. J Neurosci 30, 4590–4600. 10.1523/JNEUROSCI.0640-10.2010.

73. Mathys, H., Peng, Z., Boix, C.A., Victor, M.B., Leary, N., Babu, S., Abdelhady, G., Jiang, X., Ng, A.P., Ghafari, K., et al. (2023). Single-cell atlas reveals correlates of high cognitive function, dementia, and resilience to Alzheimer’s disease pathology. Cell 186, 4365–4385.e27. 10.1016/J.CELL.2023.08.039.

74. Kim, D.K., Han, D., Park, J., Choi, H., Park, J.C., Cha, M.Y., Woo, J., Byun, M.S., Lee, D.Y., Kim, Y., et al. (2019). Deep proteome profiling of the hippocampus in the 5XFAD mouse model reveals biological process alterations and a novel biomarker of Alzheimer’s disease. Exp Mol Med 51. 10.1038/S12276-019-0326-Z.

75. Johnson, E.C.B., Dammer, E.B., Duong, D.M., Ping, L., Zhou, M., Yin, L., Higginbotham, L.A., Guajardo, A., White, B., Troncoso, J.C., et al. (2020). Large-scale proteomic analysis of Alzheimer’s disease brain and cerebrospinal fluid reveals early changes in energy metabolism associated with microglia and astrocyte activation. Nat Med 26, 769–780. 10.1038/S41591-020-0815-6.

76. Mulder, J., Zilberter, M., Pasquaré, S.J., Alpár, A., Schulte, G., Ferreira, S.G., Köfalvi, A., Martin-Moreno, A.M., Keimpema, E., Tanila, H., et al. (2011). Molecular reorganization of endocannabinoid signalling in Alzheimer’s disease. Brain 134, 1041–1060. 10.1093/BRAIN/AWR046.

77. Burrinha, T., Martinsson, I., Gomes, R., Terrasso, A.P., Gouras, G.K., and Almeida, C.G. (2021). Upregulation of APP endocytosis by neuronal aging drives amyloid-dependent synapse loss. J Cell Sci 134. 10.1242/JCS.255752.

78. Sun, N., Youle, R.J., and Finkel, T. (2016). The Mitochondrial Basis of Aging. Mol Cell 61, 654. 10.1016/J.MOLCEL.2016.01.028.

79. Singh, P.B., and Zhakupova, A. (2022). Age reprogramming: cell rejuvenation by partial reprogramming. Development 149. 10.1242/DEV.200755.

80. López-Otin, C., Blasco, M.A., Partridge, L., Serrano, M., and Kroemer, G. (2023). Hallmarks of aging: An expanding universe. Cell 186, 243–278. 10.1016/J.CELL.2022.11.001.

81. Montalban, E., Al-Massadi, O., Sancho-Balsells, A., Brito, V., de Pins, B., Alberch, J., Ginés, S., Girault, J.A., and Giralt, A. (2019). Pyk2 in the amygdala modulates chronic stress sequelae via PSD-95-related micro-structural changes. Transl Psychiatry 9, 1–12. 10.1038/s41398-018-0352-y.

82. Giralt, A., de Pins, B., Cifuentes-Díaz, C., López-Molina, L., Farah, A.T., Tible, M., Deramecourt, V., Arold, S.T., Ginés, S., Hugon, J., et al. (2018). PTK2B/Pyk2 overexpression improves a mouse model of Alzheimer’s disease. Exp Neurol 307, 62–73. 10.1016/J.EXPNEUROL.2018.05.020.

83. Pérez-Sisqués, L., Sancho-Balsells, A., Solana-Balaguer, J., Campoy-Campos, G., Vives-Isern, M., Soler-Palazón, F., Anglada-Huguet, M., López-Toledano, M.-Á., Mandelkow, E.-M., Alberch, J., et al. (2021). RTP801/REDD1 contributes to neuroinflammation severity and memory impairments in Alzheimer’s disease. Cell Death Dis 12, 616. 10.1038/s41419-021-03899-y.

84. Tronche, F., Kellendonk, C., Kretz, O., Gass, P., Anlag, K., Orban, P.C., Bock, R., Klein, R., and Schütz, G. (1999). Disruption of the glucocorticoid receptor gene in the nervous system results in reduced anxiety. Nat Genet 23, 99–103. 10.1038/12703.

85. Giralt, A., Brito, V., Chevy, Q., Simonnet, C., Otsu, Y., Cifuentes-Díaz, C., de Pins, B., Coura, R., Alberch, J., Ginés, S., et al. (2017). Pyk2 modulates hippocampal excitatory synapses and contributes to cognitive deficits in a Huntington’s disease model. Nat Commun 8, 15592. 10.1038/ncomms15592.

86. Schneider, C.A., Rasband, W.S., and Eliceiri, K.W. (2012). NIH Image to ImageJ: 25 years of image analysis. Nat Methods 9, 671–675. 10.1038/nmeth.2089.

87. Carpenter, A.E., Jones, T.R., Lamprecht, M.R., Clarke, C., Kang, I.H., Friman, O., Guertin, D.A., Chang, J.H., Lindquist, R.A., Moffat, J., et al. (2006). CellProfiler: Image analysis software for identifying and quantifying cell phenotypes. Genome Biol 7, 1–11. 10.1186/GB-2006-7-10-R100/FIGURES/4.

88. Susaki, E.A., Tainaka, K., Perrin, D., Kishino, F., Tawara, T., Watanabe, T.M., Yokoyama, C., Onoe, H., Eguchi, M., Yamaguchi, S., et al. (2014). Whole-brain imaging with single-cell resolution using chemical cocktails and computational analysis. Cell 157, 726–739. 10.1016/j.cell.2014.03.042.

89. Yang, B., Treweek, J.B., Kulkarni, R.P., Deverman, B.E., Chen, C.K., Lubeck, E., Shah, S., Cai, L., and Gradinaru, V. (2014). Single-cell phenotyping within transparent intact tissue through whole-body clearing. Cell 158, 945–958. 10.1016/J.CELL.2014.07.017.

